# GWAs reveals SUBER GENE1-mediated suberization via Type One Phosphatases

**DOI:** 10.1101/2025.05.06.652434

**Authors:** Jian-Pu Han, Linnka Lefebvre-Legendre, Jun Yu, Maria Beatriz Capitão, Chloé Beaulieu, Kay Gully, Vinay Shukla, Yibo Wu, Andreas Boland, Christiane Nawrath, Marie Barberon

## Abstract

Suberin deposition in the root endodermis is critical for plant nutrient acquisition and environmental adaptation. Here, we used an unbiased forward genetic approach based on natural variation across 284 *Arabidopsis thaliana* accessions to identify novel regulators of endodermal suberization. This screen revealed striking diversity in suberin levels and patterns, uncovering broader roles for suberin beyond those observed in the reference accession Col-0. A genome-wide association study pinpointed *SUBER GENE1* (*SBG1*), a previously uncharacterized gene encoding a 129-amino acid protein, as a key regulator of suberin deposition. SBG1 acts through physical interaction with type one protein phosphatases (TOPPs) via conserved SILK and RVxF motifs. Disrupting this interaction abolishes SBG1 function, while *topp* mutants exhibit enhanced endodermal suberization, mirroring *SBG1* overexpression. Our findings uncover a previously unknown regulatory module linking suberin formation to TOPP activity and ABA signaling and provide a framework for improving plant stress resilience through targeted manipulation of root barrier properties.

## Main

Plant roots serve as the primary interface between plants and the soil, playing a critical role in nutrient acquisition and water uptake. To regulate the movement of water, ions, and nutrients, roots have evolved complex anatomical structures, including the deposition of suberin in the root endodermis. Suberin, a hydrophobic biopolymer, forms a selective barrier that not only controls radial transport but also protects plants from environmental stresses such as drought, salinity, nutrient imbalances, and pathogens^1^. The pattern of suberin deposition in the root endodermis is precisely regulated during root development^1^. In *Arabidopsis thaliana*, endodermal differentiation progresses through distinct stages. Initially, Casparian strips (CS) form an apoplastic barrier (non-suberized region) in the endodermis. Following this, patches of endodermal cells undergo suberization (patchy suberization zone), where suberin is selectively deposited. Eventually, the entire endodermal layer becomes uniformly suberized (suberized zone) (Fig. 1a). At the molecular level, the pattern of endodermal suberization is a highly regulated process, orchestrated by the interplay of several hormones and signaling pathways^1^. Among them abscisic acid (ABA) plays a central role in both developmental and stress-induced suberin formation^2^. Exogenous ABA treatments strongly upregulate the expression of genes involved in suberin biosynthesis, monomer transport and polymerization and lead to ectopic endodermal suberization close to the root tip^2–7^. Conversely, ABA-insensitive mutants, including lines expressing the dominant-negative *abi1-1* allele of *ABA INSENSITIVE1* specifically in the endodermis, exhibit markedly reduced suberin deposition^2,3,8,9^. These findings demonstrate that endodermal ABA signaling is not only essential for stress-induced suberization but also required for proper developmental establishment of the endodermal suberin barrier. Additionally, CIF peptides (CASPARIAN STRIP INTEGRITY FACTOR), interacting with the Leucine-Rich Repeat Receptor-Like Kinase SGN3/GSO1 (SCHENGEN3/GASSHO1), regulate the integrity and proper function of CS by reinforcing endodermal barriers, including suberization^10,11^ independently of ABA^3,8^. Recent studies have also highlighted MYB (MYELOBLASTOSIS) transcription factors, acting downstream of ABA, such as MYB9, MYB39, MYB41, MYB53, MYB92, and others^3,7,12,13^, as key regulators controlling suberin biosynthesis, transport, and polymerization. The complexity of this regulatory network underscores suberin’s central role in stress adaptation and environmental acclimation. Despite substantial progress in identifying the molecular components involved in suberin biosynthesis and regulation, no unbiased forward genetic screen has yet been conducted to discover regulators of endodermal suberization. Furthermore, natural variation in suberin deposition across plant species and genotypes remains largely unexplored.

**Fig. 1.**
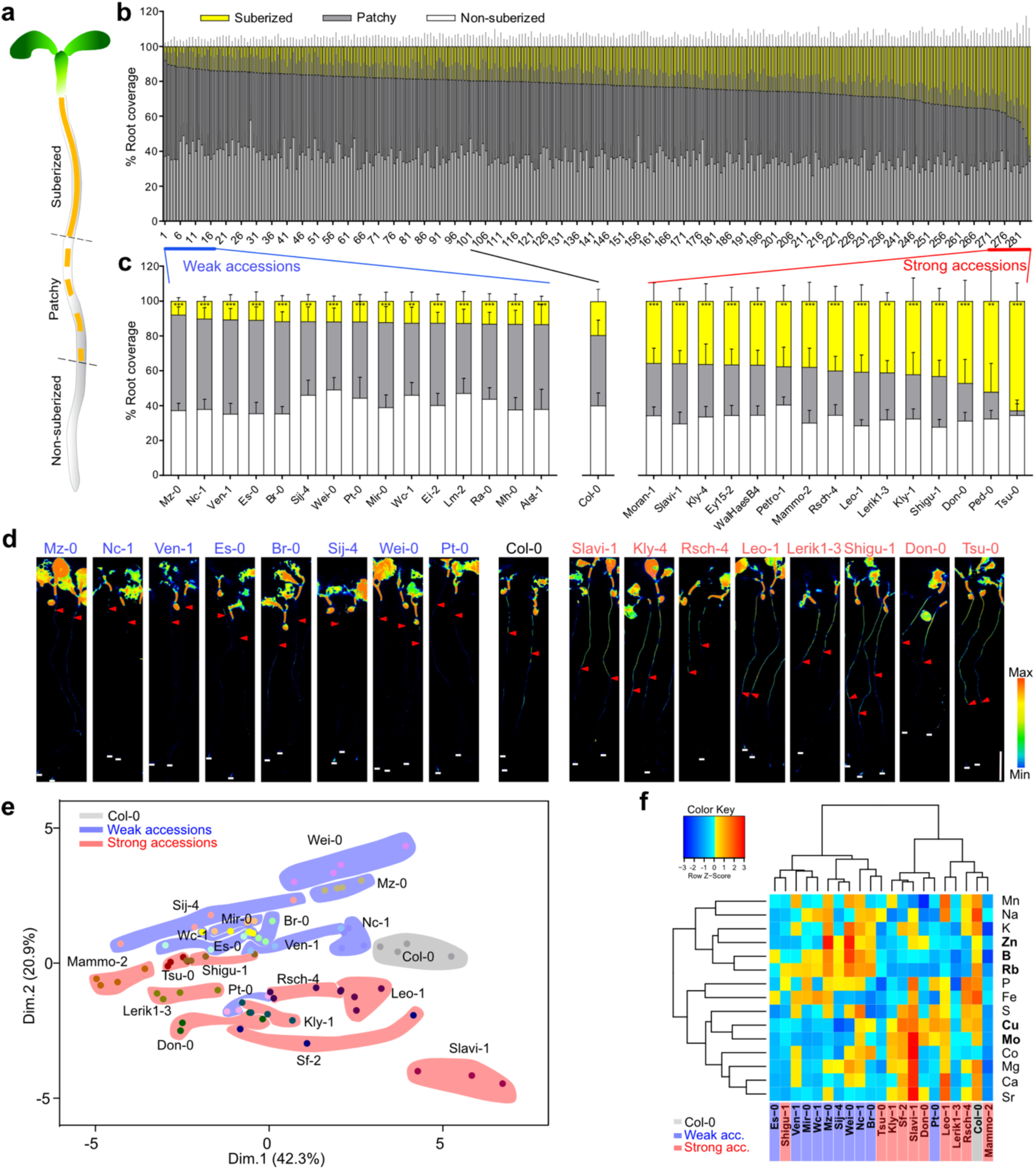
Natural variation in suberin deposition among *Arabidopsis* accessions. **a**, Schematic view of the three zones of suberization in roots. **b**, Quantification of suberin deposition in 4-day-old roots of 284 *Arabidopsis* accessions using Fluorol Yellow staining. Accessions are sorted by the percentage of suberized zone. Data are presented as stacked columns showing the mean percentage of each suberization zone + s.d., *n* ≥ 10. Numeric values are provided in Supplementary Table 1. **c**, Comparison of suberin deposition in Col-0 and selected accessions representing the extremes (weak and strong) of the distribution shown in (**b**). Statistical differences in the suberized zone were assessed by one-way ANOVA followed by Dunnett’s test and are presented for the comparison to Col-0 (\*\**P* < 0.01, \*\*\**P* < 0.001). **d**, Whole-mount images of Fluorol Yellow-stained roots from selected extreme accessions. Red arrows indicate the onset of the suberized zone; white lines mark the root tip. Scale bar, 2 mm. **e**, Principal component analysis (PCA) of ionomic profiles in Col-0 and 20 extreme accessions. Accessions with weak and strong suberization are indicated with blue and red areas respectively (hand drawn). *n* = 4. **f**, Hierarchical clustering of ionomic profiles from the same set of extreme accessions. The 5 elements contributing most to the clustering are highlighted in bold. Data for (**e**-**f**) are available in Supplementary Table 3.

In this study, we investigated natural variation in suberin deposition within the root endodermis. By analyzing a set of 284 *A. thaliana* accessions, we identified natural variants with distinct suberin deposition patterns and demonstrated that suberin plays a central role in mineral accumulation, extending beyond its previously established functions in the reference ecotype Col-0. Through a genome-wide association study (GWAS), we identified *SUBER GENE 1 (SBG1*), a previously uncharacterized gene encoding a 129-amino-acids protein that plays a key role in suberin regulation. We further discovered that SBG1 interacts with Type One Protein Phosphatases (TOPPs) via conserved SILK and RVxF motifs and inhibits the phosphatase activity positioning it as a novel TOPPs regulator involved in fine-tuning endodermal suberization. These findings reveal a previously unknown regulatory mechanism governing suberin formation and provide new insights into the genetic determinants of suberization and its role in plant adaptation to environmental conditions.

## Results

### Natural variation for endodermal suberin

To explore the variation of endodermal suberin deposition in *A. thaliana*, we analyzed the suberin pattern in 284 genetically diverse natural accessions. We stained 4-day-old seedlings with Fluorol Yellow to visualize suberin and quantified endodermal suberization by measuring the root segments that were non-suberized, patchy, or fully suberized. A wide range of suberin patterns was observed, particularly in the fully suberized zone (Fig. 1b, Supplementary Table 1). For further analysis, we selected 15 accessions with a significantly reduced proportion of fully suberized cells compared to Col-0 (hereafter referred to as “weak” accessions) and 15 accessions with a significantly extended fully suberized zone (hereafter referred to as “strong” accessions), as determined by quantitative analysis of suberin deposition patterns (Fig. 1c,d). Root length is known to vary across accessions^14,15^, but we found no significant correlation between root length and the fully suberized zone and only weak correlations with the non-suberized and patchy zones (Extended Data Fig. 1a,b, Supplementary Table 1). Minor meristem size variations in specific accessions (e.g., Wei-0, Pt-0, and Lerik1-3) did not correlate with suberin patterns (Extended Data Fig. 1c). The above indicates that the observed differences in suberin pattern are not simply a consequence of differences in root growth rate. Since defects in CS formation are known to cause enhanced suberization^11,16–20^, we additionally measured CS functionality using the apoplastic tracer propidium iodide (PI). We confirmed that CS functionality remained intact across the extreme accessions, ruling out defective CS formation as a driver for early or ectopic suberization (Extended Data Fig. 1d). Taken together, these data point to genetic variation directly affecting endodermal suberization, potentially linked to adaptive responses. The geographical distribution of all accessions, particularly those with extreme suberin deposition levels, was mapped, revealing a trend with strong accessions predominantly located in southern Europe, the Mediterranean region, and continental Eurasia (Extended Data Fig. 2a). To further investigate potential environmental influences, we analyzed correlations between suberin deposition and 473 climate variables (Supplementary Table 2). To account for the population structure, the genotype matrix of accessions was summarized by PCA and the first 2 PCs (principal components) were taken as covariates in the linear model for correlation analysis. Several strong positive correlations were identified, including those with temperature during the warmest month, summer solar radiation, precipitation seasonality, and the global terrestrial drought index (Extended data Fig. 2b-d). Consistent with these correlations, endodermal suberization in Col-0 increased following 24-hour exposure to elevated temperatures (from 22°C to 30°C) or osmotic stress induced by mannitol treatment, which mimics reduced water availability (Extended data Fig. 2e,f). Notably, Tsu-0, one of the accessions displaying enhanced suberin deposition in our screen, was reported to be drought tolerant^21^, a trait that may be partly attributable to its increased suberization. Together, these findings are supporting the idea that the observed suberin variations are adaptive responses to local environmental factors, particularly temperature and water availability.

### Suberin variation is associated with specific mineral imbalance

We leveraged the extreme accessions to investigate suberin function in nutrient homeostasis, a process extensively studied in Col-0^2,3,8,13^. Given that in Col-0, salt stress induces endodermal suberization and that suberin acts as barrier to limit sodium (Na) entry^2,3,22,23^, we first assessed suberin induction under moderate salt stress conditions, under which Col-0 displays only a mild response (Extended Data Fig. 3a-d). In this context, all weak accessions showed an increase in suberin, though to varying degrees. Notably, Sij-4, Wei-0, and Mir-0 exhibited minimal induction, indicating a reduced sensitivity to salt-triggered suberization (Extended Data Fig. 3a,b). In contrast, most strong accessions, except Slavi-1, became fully suberized, with Shigu-1, Tsu-0, and Don-0 showing a strong reduction of the non-suberized zone, consistent with a hypersensitivity response (Extended Data Fig. 3c,d). Growth assays further revealed that several weak accessions maintained shoot biomass under salt stress, whereas most strong accessions displayed severe growth reduction (Extended Data Fig. 3e-g). To compare the magnitude of salt effects across accessions, we calculated Cohen’s *d* for the change in shoot biomass between control and salt conditions. All accessions exhibited negative effect sizes (|*d*| = 0.61–3.72), reflecting reduced biomass upon salt treatment. The magnitude of this reduction varied substantially among genotypes: Col-0 showed an intermediate effect (|*d*| = −1.41), similar to the weak accessions Nc-1, Es-0 and Pt-0 and the strong accessions Leo-1, Don-0 and Tsu-0 ((|*d*| = 1.11–1.99). The smallest effects (|*d*| = 0.61–0.81) were observed in three weak accessions (Wei-0, Sij-4 and Br-0), whereas the strongest effects (|*d*| = 3.24–3.72) were restricted to one weak accession (Mz-0) and three strong accessions (Mammo-2, Sf-2 and Kly-1). These patterns reveal accession-specific differences in salt sensitivity with a partial separation between weak and strong suberizing accessions. These results contrast with the established model from Col-0 in which reduced suberin increases salt sensitivity and enhanced suberin confers protection^2,8^. To test this further, we expressed *CUTICLE DESTRUCTING FACTOR1* (*CDEF1*) under the *ELTP* promoter to specifically degrade endodermal suberin in strong accessions (Extended Data Fig. 4a), as previously demonstrated in Col-0^2,24,25^, and examined their response to salt stress in plates and soil (Extended Data Fig. 4b,c). Consistent with prior reports *ELTP::CDEF1* expression in Col-0 increased salt sensitivity, as shown by a stronger reduction in shoot biomass in both growth conditions and a light-brown discoloration. A similar effect was observed in the strong accession Slavi-1, which already differed from other strong accessions in its suberin response to salt. In contrast, suberin degradation in all other strong accessions tested, did not alter their salt sensitivity on plates and only lead to a mild growth reduction and light-brown discoloration in soil, further supporting that suberin functions in an accession-dependent manner in salt stress responses, distinct from the model established in Col-0.

To further explore the link between suberin variation and mineral homeostasis beyond Na, we conducted ionomic profiling of shoots from 20 extreme accessions alongside Col-0 grown on agar plates. Principal component and hierarchical clustering revealed distinct ionomic signatures between weak and strong accessions, particularly for zinc (Zn), boron (B), rubidium (Rb), copper (Cu) and molybdenum (Mo), but not Na (Fig. 1e,f, Extended Data Fig. 4d,e, Supplementary Table 3). These results highlight the pivotal role of endodermal suberization in mineral acquisition and confirm trends observed in suberin-altered Col-0 mutants and lines in similar growth conditions. Notably, B was previously shown to accumulate at lower levels in Col-0 plants with ectopic endodermal suberin and at higher levels in those with reduced suberin^3,18^. Similarly, Rb levels were reduced, while Cu and Mo levels increased in Col-0 plants with enhanced suberization^3,13,18^. However, our data also reveal differences from prior findings: Na, which typically accumulates in suberin-deficient Col-0 plants, did not vary significantly here, while Zn, lower in strong accessions in our study, has not previously been identified as a hallmark of suberin-dependent ionomic shifts. We therefore examined whether the lower Zn content observed in strong accessions might be accompanied by altered physiological responses to Zn deficiency. Under Zn-limiting conditions, the strong accessions Sf-2 and Leo-1, as well as the weak accession Wei-0, displayed a pronounced reduction in shoot biomass (Extended Data Fig. 5a,b). To compare the magnitude of Zn deficiency effects among accessions, we calculated Cohen’s *d* for the change in shoot biomass between control and –Zn conditions. All accessions exhibited large negative effect sizes (|*d*| = 3.75–8.28), consistent with a strong reduction in growth under Zn limitation. However, the effect size varied between genotypes, with Wei-0 and Leo-1 showing the largest reduction (|*d*| < −7) and Col-0 the smallest (|*d*| = −3.75), revealing accession-specific differences in Zn sensitivity without a clear separation between the weak and strong accessions analyzed. In addition, the strong accessions Sf-2 and Leo-1 developed characteristic Zn-deficiency symptoms in leaves, a phenotype not observed in Col-0 or in the weak accessions Mz-0 and Wei-0 (Extended Data Fig. 5a,b). To clarify whether suberin contributes to these differences, we expressed *ELTP::CDEF1* to degrade suberin in several strong accessions, as described above for salt stress (Extended Data Fig. 4a, Extended Data Fig. 5c-e). In Col-0, suberin degradation had no effect on shoot biomass under Zn limitation but mitigated chlorophyll loss compared to WT. No significant effect of *ELTP::CDEF1* expression was observed in Kly-1 and Rsch-4. However, in Tsu-0 and Shigu-1, suberin degradation alleviates the negative effect of Zn limitation on both shoot biomass and chlorosis. Together, these results demonstrate an accession-dependent role of endodermal suberin in Zn, neutral in some accessions but exacerbating Zn sensitivity in others. They reveal a previously underappreciated role of endodermal suberization in Zn acquisition and highlight a trade-off between protective barrier function and micronutrient uptake. While they challenge the view of suberin as a strict protective barrier against Na, our findings underscore its impact on Zn homeostasis, revealing increased suberin deposition correlates with reduced Zn accumulation and heightened sensitivity to Zn limitation in specific genetic backgrounds.

### GWAS for endodermal suberin deposition identifies *SUBER GENE 1* (*SBG1*)

To uncover the genetic basis of natural variation in endodermal suberin deposition, we performed a Genome Wide Association Study (GWAS) for the percentage of the fully suberized zone (Fig. 1b, Extended Data Fig. 6a-c, Supplementary Table 1)^26^. The pseudo-heritability (h^2^) for the suberized zone was estimated to be 0.71 in GWA-Portal, indicating a substantial genetic contribution to this trait. GWAs identified two significant loci on chromosomes 1 and 4 (Fig. 2a) with the lead SNP (*G*/*T*) on chromosome 1 located 319 bp downstream of *AT1G52565*, a previously uncharacterized gene encoding a 129 aa protein, later designated SUBER GENE1 (SBG1) (Fig. 2b). On Chromosome 4, two SNPs mapped between *AT4G05475* and *AT4G05490* encoding putative F-BOX proteins but given their greater distance from the SNPs (1839 bp and 1236 bp, respectively) (Extended Data Fig. 6d), we focused on the Chromosome 1 locus. Accessions carrying the *G* allele at this locus exhibited significantly higher suberin levels than those with the *T* allele (Extended Data Fig. 6e).

**Fig. 2.**
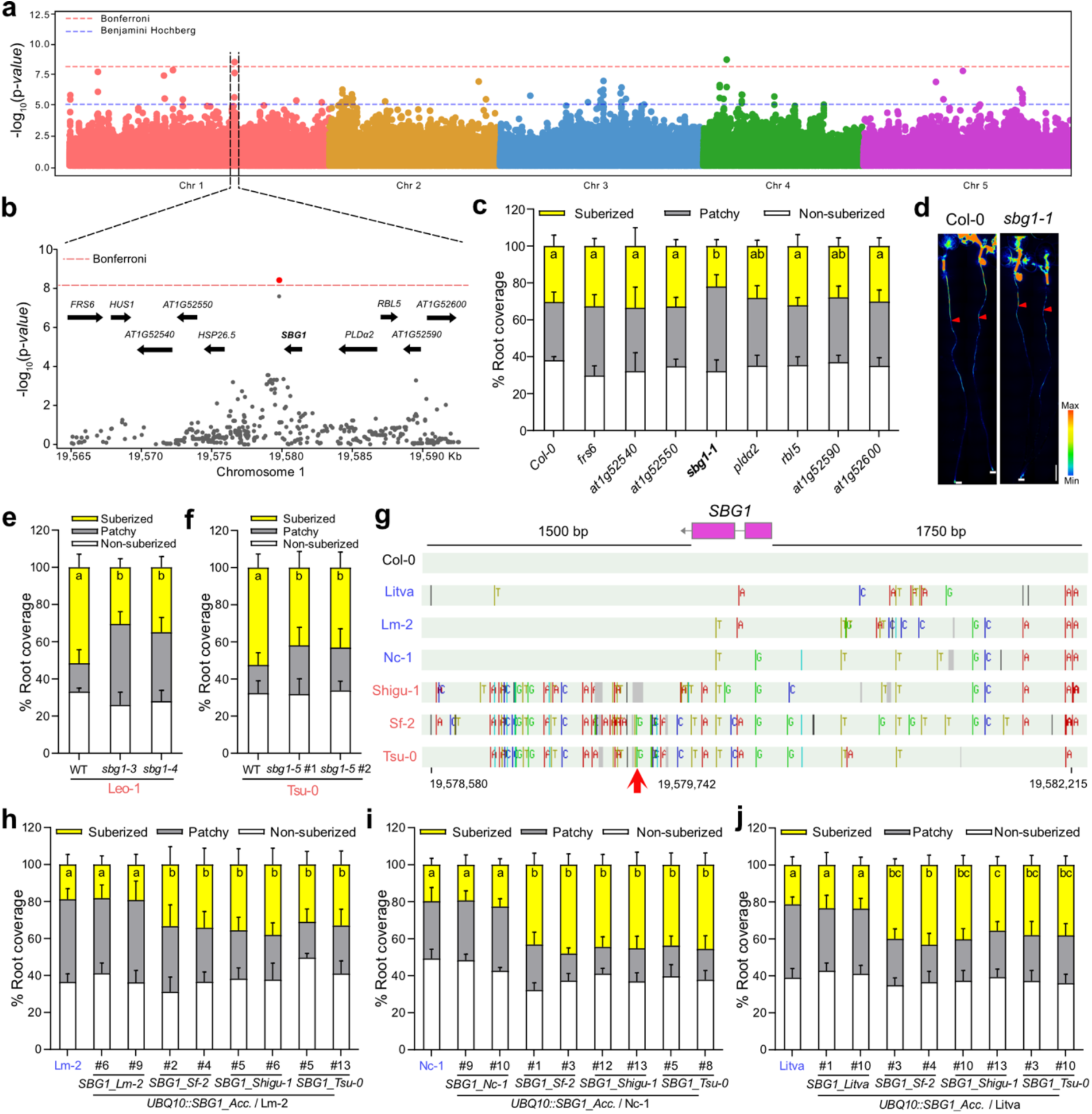
Identification of *SBG1* as a genetic determinant of suberin variations. **a**, Manhattan plot of GWAS for suberin variations across 265 accessions. Horizontal dashed lines indicate significant thresholds after multiple testing correction: red for Bonferroni and blue for Benjamini–Hochberg (FDR = 10%). **b**, SNPs and annotated genes surrounding the *SBG1* genomic region. Each dot represents a SNP, with the top SNP (19,579,742 bp) highlighted in red. **c**, Suberin deposition pattern in T-DNA insertion lines for candidate genes shown in (**b**). Data are presented as stacked columns showing the mean percentage of each suberization zone + s.d., *n* = 10. One-way ANOVA followed by a Tukey’s test was used to assess significance for the suberized zone; different letters indicate significant differences between genotypes (*P* < 0.05). **d**, Whole-mount images of Fluorol Yellow-stained roots from Col-0 and *sbg1-1* shown in (**c**). Red arrows indicate the onset of suberized zone; white lines mark the root tip. Scale bar, 1 mm. **e**,**f**, Suberin deposition pattern in *sbg1* mutants generated in Leo-1 (**e**) and Tsu-0 (**f**) backgrounds. Data are presented as in (**c**), *n* = 10. One-way ANOVA followed by a Tukey’s test was used to test significance in the suberized zone (*P* < 0.05). **g**, SNP profiles in the *SBG1* genomic region across 3 *T* and 3 *G* allele accessions. The top SNP from GWAs is marked with a red arrow. SNP data were obtained from the 1001 Genomes dataset. **h**-**j**, Suberin deposition pattern in transgenic lines expressing *SBG1* under the *UBQ10* promoter in the background of Lm-2 (**h**), Nc-1 (**i**) and Litva (**j**). *SBG1* genomic sequence including 1.5 kb downstream of the stop codon was derived from Lm-2, Nc-1, Litva, Sf-2, Shigu-1 or Tsu-0 as indicated. Two independent lines were analyzed per construct. Data are presented as in (**c**). One-way ANOVA followed by a Tukey’s test was used to determine significance in the suberized zone (*P* < 0.05).

To validate the role of *SBG1*, we quantified suberin deposition in available T-DNA insertion lines in Col-0 for genes within ±15 Kb of the lead SNP (Fig. 2c). Only the *sbg1-1* mutant showed significantly reduced suberin compared to WT (Fig. 2c,d, Extended Data Fig. 6f). This phenotype was independently confirmed in CRISPR-Cas9 generated *sbg1* mutants in the strong accessions Leo-1 and Tsu-0, further supporting the role of *SBG1* in suberization (Fig. 2e,f, Extended Data Fig. 6h,I). To explore how allelic variation influences suberization we compared *SBG1* sequences between weak (*T* alleles) and strong (*G* alleles) accessions. Protein alignment revealed a four-amino acid insertion (REQR in place of S83) in the strong alleles, but structural prediction using AlphaFold did not indicate any effect on protein folding allowing to formulate hypothesis on a functional effect of this insertion (Extended Data Fig. 7a,b). Sequence comparison of *T* and *G* alleles revealed several additional SNPs clustered in the 3’ region (Fig. 2g). Because 3’ UTR often regulate mRNA stability and translation, we hypothesized that these polymorphisms could affect *SBG1* expression. However, qRT-PCR analysis did not reveal a consistent correlation between *SBG1* transcript levels and suberization across accessions (Extended Data Fig. 7c). Interestingly, secondary structure predictions for the 3’UTR showed pronounced differences between *T* and *G* alleles (Extended Data Fig. 7d,e), suggesting potential regulatory effects that merit further investigation. To functionally assess allele-specific effects, we cloned *SBG1* genomic sequences, including 1.5 kb of the 3’ region, from accessions carrying either the *G* allele (strong accessions, Shigu-1, Sf-2 and Tsu-0) or the *T* allele (weak accessions Lm-2, Nc-1 and Litva). When expressed under the constitutive and ubiquitous *UBQ10* promoter in *T* allele backgrounds (Lm-2, Nc-1 and Litva), *T* allele transgene did not alter suberization, whereas *G* allele constructs significantly enhanced suberin deposition near the root tip (Fig. 2h-j). Together, these findings establish *SBG1* as a key positive regulator of suberization across natural accessions, demonstrating that allelic variations in *SBG1* modulate suberin deposition in the root endodermis.

### *SBG1* is a positive regulator of endodermal suberization

To further investigate the role of SBG1 in suberin deposition, we functionally characterized this gene in Col-0. The reduced suberin phenotype observed in *sbg1-1* (Fig. 2c,d) was confirmed with a second allele, *sbg1-2*, generated by CRISPR-Cas9 (Fig. 3a,b, Extended Data Fig. 8a). Complementation of *sbg1-1* with either SBG1 or SBG1-mVenus under its native promoter fully restored suberization (Fig. 3a,b). Conversely, constitutive expression of SBG1-mVenus under the *UBQ10* promoter significantly enhanced suberization near the root tip (Fig. 3c,d), consistent with the enhances suberization observed when *G* alleles were expressed in weak accessions (Fig. 2h-j). Although no significant difference in total suberin content was detected in *sbg1-1* mutants compared to Col-0, the enhanced suberization in *UBQ10::SBG1-mVenus* lines was corroborated by chemical analysis of whole root suberin content, revealing a significant increase in suberin monomers, with a 34% increase in total aliphatic content (Fig. 3e.f). Noteworthy, the most abundant monomers were particularly increased, C22:0 and C24:0 fatty acids, C18:1 dicarboxylic acid (DCA), and C18:1 ω-hydroxy fatty acid, corresponding to the main monomers also increased after ABA application^2^ or after *MYB41* endodermal overexpression^3^. Together, these results demonstrate that *SBG1* is both necessary and sufficient to promote endodermal suberin deposition.

**Fig. 3.**
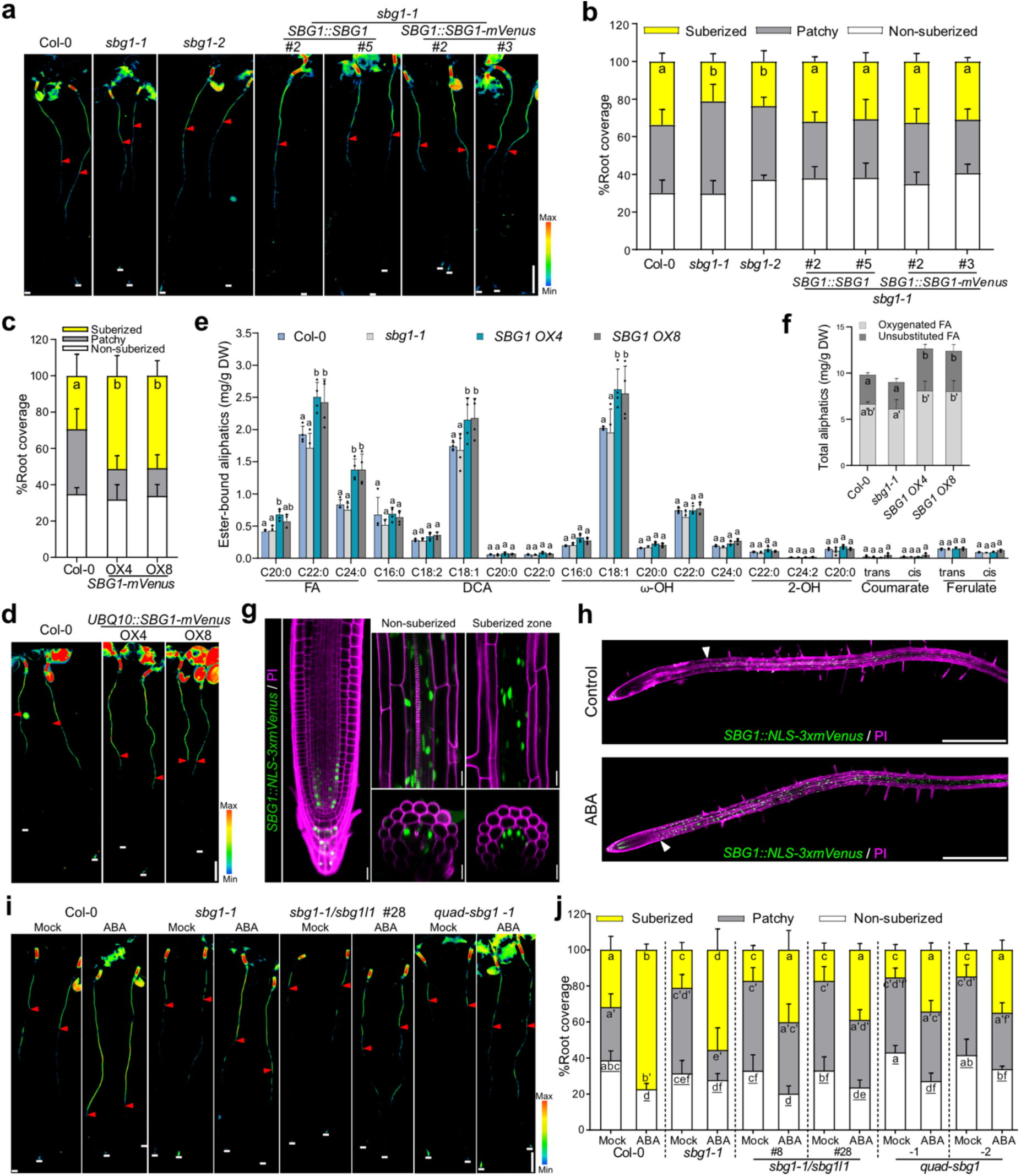
*SBG1* regulates suberization. **a,b**, Whole-mount images of Fluorol Yellow-stained roots (**a**) and suberin deposition pattern (**b**) in *sbg1* mutants and complemented lines. Red arrows indicate the onset of suberized zone; white lines mark the root tip. Scale bar, 1 mm. Data are presented as stacked columns showing the mean percentage of each suberization zone + s.d., *n* = 10. One-way ANOVA followed by a Tukey’s test was used to assess significance for the suberized zone; different letters indicate significant differences between genotypes (*P* < 0.05). **c**,**d**, Whole-mount images (**c**) and quantification (**d**) of suberin deposition in transgenic lines expressing *SBG1-mVenus* under the *UBQ10* promoter. Data are presented as in (**a,b**). **e**,**f**, Quantification of ester-bound aliphatic and aromatic suberin monomers in 5-day-old roots. FA, fatty acid; DCA, dicarboxylic fatty acid; ω-OH, ω-hydroxy fatty acid; DW, dry weight. Data are shown as histograms with individual values overlaid, s.d., *n* = 4 pools of 200-300 roots. Two-way ANOVA followed by a Tukey’s test was used to assess significance for each suberin monomer, oxygenated and unsubstituted FA; different letters indicate significant differences between genotypes (*P <* 0.05). **g**, Confocal images of *SBG1::NLS-3xmVenus* line showing mVenus signal (green) combined with PI staining (magenta) to highlight cell walls. In the non-suberized and suberized root zones, mVenus signal was projected in 3D. Cross-section views were generated by re-slicing Z-stacks. Scale bar, 20 µm. **h**, Confocal images of *SBG1::NLS-3xmVenus* lines treated with or without 1 µM ABA for 18h. mVenus signal is shown in green and PI in magenta. White arrows indicate the onset of mVenus signal after the division zone. Scale bar, 500 µm. **i**,**j**, Whole-mount images (**i**) and suberin deposition pattern (**j**) in *sbg1-1* and high-order *sbg1l* mutants under control (mock) or 1 µM ABA treatment for 18h. 4-day-old seedlings were transferred to either ½ MS plates or ½ MS plate + ABA. Data are presented as in (**a,b**). Different letters indicate significant differences in the different suberized zones between genotypes and conditions with apostrophes and underlines for the patch and non-suberized zones respectively.

A defining feature of genes involved in endodermal suberin formation is their specific expression in this tissue and their induction by ABA^2,3,6,7,12,27^. Whole-root expression analysis revealed that *SBG1* was moderately upregulated after ABA treatment (Extended Data Fig. 8b), reaching levels comparable to *CYP86A1* (also known as *HORST, HYDROXYLASE OF ROOT SUBERIZED TISSUE*), a cognate suberin biosynthetic enzyme^28^. Notably, this induction is markedly weaker than that of *MYB41,* a major transcriptional activator of endodermal suberization^3^. Such a moderate increase, while clearly ABA-responsive, might reflect a function in suberin biosynthesis or as a fine-tuning component rather than a primary regulator within the endodermal suberin network. To examine *SBG1* expression *in situ*, we generated *SBG1::NLS-3xmVenus* transcriptional reporter lines. Promoter activity was initially detected in the columella and early endodermal lineage cells at the root tip (Fig. 3g). In differentiated roots, *SBG1* promoter activity became predominantly endodermal, with weaker activity observed in vascular tissues (Fig. 3g). Following ABA treatment, promoter activity expanded into the early pericycle and endodermal cell lineages and was additionally detected in some epidermal and cortical cells in the elongation and early differentiation zones (Fig. 3h, Extended Data Fig. 8c). Moreover, consistent with suberin-related genes such as *MYB41, MYB92,* and *GPAT5*^2,3^, ABA further enhanced *SBG1* promoter activity in the endodermis, extending expression toward the meristematic zone (Fig. 3h, Extended Data Fig. 8c).

The *A. thaliana* genome contains three uncharacterized homologs of *SBG1*, hereafter referred to as *SBG1-Like1, SBG1-Like2* and *SBG1-Like3* (*SBG1L1-3*) (Extended Data Fig. 8d). Under control conditions, the double mutant *sbg1;sbg1l1* was phenotypically similar to *sbg1*, while the quadruple mutant *sbg1;sbg1l1;sbg1l2;sbg1l3* (*quad-sbg1*) displayed only a slightly extended non-suberized zone (Fig. 3i,j). Under ABA treatment, however suberin induction was progressively attenuated from *sbg1* to *sbg1;sbg1l1,* and most strongly in *quad-sbg1*, which exhibited a suberization pattern resembling that of untreated Col-0 roots. These results suggest that SBG1 and its homologs contribute additively but modestly to ABA-mediated suberization response in endodermal cells.

To test this hypothesis, we compared the transcriptional responses of Col-0 and *sbg1* roots to ABA by RNA-seq. Given our hypothesis that *SBG1* acts as a local modulator of ABA-response in a limited number of endodermal cells, we expected only subtle global transcriptional effects. Consistent with this, under control conditions only minor expression differences were detected between *sbg1* and Col-0, with 29 genes upregulated and 45 downregulated, and no enrichment for specific biological processes (Supplementary Table 4). We next compared the ABA-induced genes in Col-0 and *sbg1* and observed striking differences between genotypes with 4’689 genes affected in Col-0 and 7’449 in *sbg1* with only 59% shared with Col-0 (Extended Data Fig. 8e, Supplementary Table 4) demonstrating that *sbg1* is affected in its ABA-response. Among the 268 genes induced by ABA in Col-0 but not in *sbg1* we could find *SBG1* as expected as well as several well-known ABA-responsive genes such as *CBF3* (*C-REPEAT BINDING FACTOR 3*), *ABF4* (*ABRE BINDING FACTOR 4*) and *LEA4-1* (*LATE EMBRYOGENESIS ABONDANT 4-1*) (Extended Data Fig. 8e). These results provide strong evidence that *sbg1* is affected in its transcriptional response to ABA.

Together, these results establish SBG1 and its homologs as positive regulators of ABA-mediated endodermal suberization and reveal that SBG1 contributes to ABA responsiveness in the root. However, given that *SBG1* encodes a small 129-amino-acid protein with no known functional domains, its precise role in ABA response or suberin formation remains elusive (Extended Data Fig. 9a).

### SBG1 promotes suberization through interference with TOPPs

We hypothesized that SBG1’s molecular function might be regulatory, mediated through protein-protein interactions. We first assessed SBG1 subcellular localization with SBG1-mVenus and SBG1-mCherry fusion proteins, which exhibited a reticulated signal colocalizing with an ER marker (Fig. 4a, Extended Data Fig. 9b). We next performed immunoprecipitation (IP) of SBG1-mVenus from plants constitutively expressing the fusion protein, followed by mass spectrometry (MS) analysis. Enriched proteins were identified by comparison with an IP control from plants expressing ER-GFP. Among the most enriched proteins (Supplementary Table 5), we identified 2 Type One Protein Phosphatases (TOPPs): TOPP2 and TOPP1 (Extended Data Fig. 9c). In total, 6 out of the 9 *A. thaliana* TOPPs (TOPP1, 2, 4, 5, 7 and 8) were significantly enriched in the SBG1 IP-MS dataset (Extended Data Fig. 9c). A Fisher’s exact test confirmed this as a highly significant enrichment (*P* = 6.3x10^-5^; odds ratio = 15.5), supporting the specificality of the association of between SBG1 and TOPP proteins. The interaction between SBG1 and TOPP2, the most strongly enriched phosphatase, was further validated by co-immunoprecipitation *in planta*, using seedlings co-expressing SBG1-mVenus and TOPP2-mScarletI (Fig. 4b, Extended Data Fig. 9d).

**Fig. 4.**
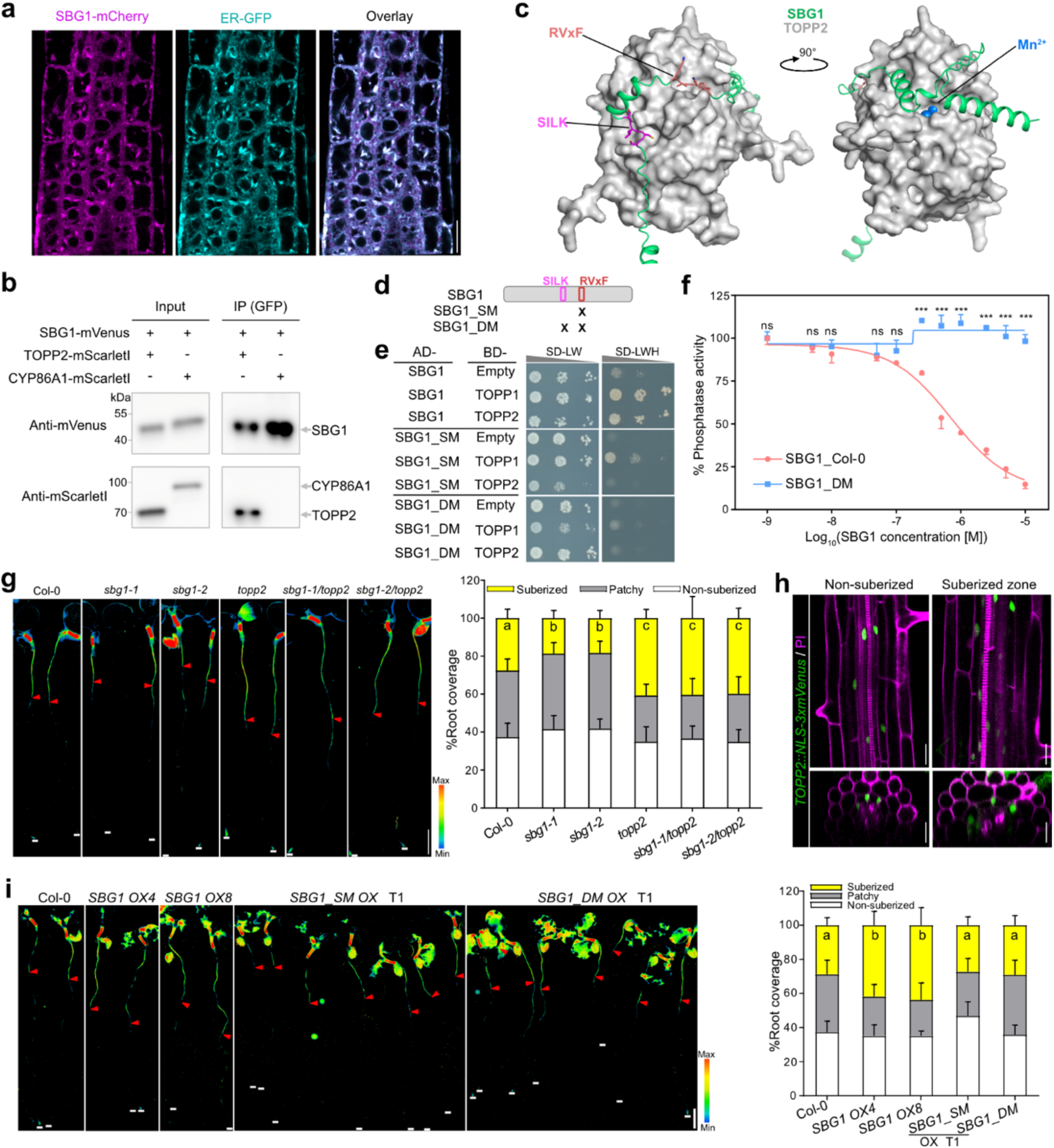
SBG1 regulates suberization through its interaction with TOPPs. **a**, Confocal images showing colocalization of SBG1- mCherry (magenta) with ER-GFP (cyan). Individual channels and merged images are shown, Scale bar, 50 µm. **b**, Co-IP assay showing SBG1 interaction with TOPP2 in roots from plants co-expressing SBG1-mVenus and TOPP2-mScarletI or SBG1-mVenus and CYP86A1-mScarletI as a negative control. See also Extended Data Fig. 9d. **c**, Predicted AlphaFold3 structure of SBG1 in complex with TOPP2. The side chains of the SILK and RVxF motifs in SBG1 and two Mn^2+^ ions in the catalytic domain of TOPP2 are highlighted. **d**, Schematic representation of mutations in the SBG1 binding motifs. SM denotes mutation in the RVxF motif, and DM denotes combined mutations in both the SILK and RVxF motifs. Mutation sites are marked with an “X”. **e**, Yeast two-hybrid assay showing interactions between WT or mutated forms of SBG1 (SM and DM) and TOPP1 or TOPP2. Drop test started with OD600 of 0.1 followed by 10 times dilutions. **f**, Phosphatase activity assay showing SBG1 concentration-dependent inhibition of TOPP2 activity *in vitro*. Two-way ANOVA followed by Sidak’s test was used to assess the significance at each concentration (\*\*\**P* < 0.001, ns, not significant, *n* = 3). **g**, Whole-mount images of Fluorol Yellow-stained roots and corresponding suberin deposition pattern in *sbg1*, *topp2* and *sbg1/topp2* double mutants (left panels). Red arrows indicate the onset of the suberized zone; white lines mark the root tip. Scale bar, 1 mm. In right panel, data are presented as stacked columns showing the mean percentage of each suberization zone + s.d., *n* = 11. One-way ANOVA followed by a Tukey’s test was used to assess significance for the suberized zone; different letters indicate significant differences between genotypes (*P* < 0.05). **h**, Confocal images of *TOPP2::NLS-3xmVenus* in roots with mVenus signal (green) overlaid with PI staining (magenta) to visualize cell walls. mVenus signal was projected in 3D. Cross-section views were generated by re-slicing Z-stacks. Scale bar, 50 µm. **i**, Whole-mount images and suberin deposition pattern in T1 overexpression lines expressing *SBG1_SM* and *SBG1_DM* compared to WT *SBG1* (*SBG1 OX4* and *8*) (left panels). In right panel, data are presented and statistically analyzed as in (**g**). *n* = 11 for Col-0 and *SBG1 OX4/*8, *n* = 20 for *SBG1_SM OX* (independent T1 lines), and *n* = 16 for *SBG1_DM OX* (independent T1 lines).

TOPPs constitute a conserved family of serine/threonine phosphatases with broad functions in plants, including cell division, differentiation, salt tolerance and ABA signaling^29^. They are orthologs of Protein Phosphatase 1 (PP1) in animals and fungi, where their activity is modulated by regulatory subunits that interact through conserved RVxF and SILK motifs^30^. These regulatory interactions can either inhibit TOPPs activity by blocking substrate access (as seen for Inhibitor-2, I2) or recruit substrates while controlling access to the catalytic site (as observed for Kinetochore Null Protein 1, KNL1)^29,31,32^. Strikingly, SBG1 and SBG1L1 both contain a SILK and RVxF motifs, while SBG1L2, and SBG1L3 retain only the RVxF motif, suggesting that SBG1 proteins might act as TOPP regulators (Extended Data Fig. 8d). To explore this hypothesis, we first employed an *in-silico* structural approach using AlphaFold3^33^, focusing on TOPP2, the most enriched interactor in our IP-MS dataset. The predicted complex showed high confidence with an interface predicted template modeling (ipTM) value of 0.85 and a predicted template modeling (pTM) of 0.79, comparable to the validated TOPP-I2 interaction^32,34^ (Fig. 4b, Extended Data Fig. 9e-g). These models predict that SBG1 interacts with TOPP2 via its central region containing both SILK and RVxF motifs (Fig. 4b, Extended Data Fig. 9e,f). Similar predictions were obtained for SBG1L1, whereas SBG1L2 and SBG1L3, which lack the SILK motif, show only partial overlap in their predicted binding interfaces (Extended Data Fig. 8d and 9g). Additionally, the C-termini of SBG1 homologs and the known TOPP inhibitor I2 are predicted to form similar structural covering over the TOPP2 catalytic site. This is particularly evident for SBG1 and SBG1L1, which possess extended C- terminal loops (Extended Data Fig. 9g), suggesting that SBG1 proteins may inhibit TOPP2 activity by sterically blocking access to its catalytic site. The differences in motif composition among SBG1 homologs and C-termini provide a mechanistic rationale for their unequal genetic contributions. SBG1 and SBG1L1, the only family members containing both the SILK and RVxF motifs required for TOPP binding and an extended C-termini, contribute most strongly to the ABA-induced suberization response, whereas SBG1L2 and SBG1L3, lacking the SILK motif, have a weaker impact. This is reflected in the incremental phenotype of the quadruple mutant compared to the double mutant (Fig. 3i), supporting a model in which SBG1 family members regulate suberization in proportion to their predicted ability to interact with TOPPs. We next experimentally validated this interaction using WT SBG1 and mutant versions carrying substitutions in the RVxF motif (SBG1_SM, V63A/F65A) or in both SILK and RVxF motifs (SBG1_DM, C46A/L47A and V63A/F65A), which were predicted to cause only a slight reduction in prediction confidence for SBG1-TOPP2 complex (Extended Data Fig. 9h). However, yeast two-hybrid assays confirmed that SBG1 interacts with both TOPP1 and TOPP2, and that the mutations SBG1_SM and SBG1_DM abolished these interactions (Fig. 4d,e). To test whether this interaction affects TOPP2 activity, we performed *in vitro* phosphatase assays using recombinant proteins. Analytic gel filtration assay showed SBG1 co-eluted with TOPP2, confirming a direct physical interaction, while the double mutant version (SBG1-DM, mutated in both motifs) failed to bind (Extended Data Fig. 10a). In phosphatase assays, SBG1 inhibited TOPP2 activity in a concentration-dependent manner, whereas SBG1-DM had no effect, demonstrating that the interaction is essential for inhibition (Fig. 4f). We next examined whether allelic variation in *SBG1* coding sequence could modulate this inhibitory effect. Comparison of the *T* and *G* alleles by AlphaFold prediction revealed no detectable differences in the predicted SBG1-TOPP2 interaction (Extended data Fig. 10b). Consistently, *in vitro* phosphatase assays using recombinant proteins from the two allelic variants showed similar TOPP2 inhibition (Extended Data Fig. 10c). These findings reinforce the hypothesis that functional differences between alleles arise from regulatory variation in the 3’UTR rather than changes from the coding sequence. Together, these findings demonstrate that SBG1 physically interacts with TOPPs, particularly TOPP2, via conserved SILK and RVxF motifs, and acts as an inhibitory regulator of their phosphatase activity.

To our knowledge, TOPPs have not previously been implicated in suberin or polyester formation. However, previous studies showed that *topp* loss-of-function mutants are hypersensitive to ABA, and TOPP1 was reported to inactivate SNF1-related protein kinase 2.6 (SnRK2.6, also known as Open stomata, OST1), a key positive regulator of ABA signalling^34,35^. Consistently, we observed that SBG1-mVenus overexpressing lines, but not *sbg1* mutants, displayed ABA hypersensitivity comparable to previously described *topp* mutants (Extended Data Fig. 11a)^35^. This suggests that SBG1 may inhibit TOPP activity, thereby modulating ABA signaling. We next examined suberin deposition in several *topp* mutants (*topp1*, *2, 6, 7, 8* and 9) and found that only *topp2* displayed a marked increase in suberin deposition, with ectopic deposition near the root tip (Fig. 4g, Extended Data Fig. 11b,c). This phenotype closely resembled that of *UBQ10::SBG1-mVenus* overexpression lines and was opposite to the reduced suberization observed in *sbg1* mutants, suggesting that SBG1 likely promotes suberization by antagonizing TOPP2 function in ABA signaling. Supporting this model, we found that *TOPP2* promoter is active in the endodermis, similar to *SBG1*, but in contrast to *SBG1*, not responsive to ABA (Fig. 4h, Extended Data Fig. 11d). This is consistent with our RNA-seq analysis showing that *TOPP2* transcript levels in roots are unaffected by ABA treatment (Extended Data Fig. 8e). To determine the genetic relationship between the two genes, we performed epistasis analysis. The double mutant *sbg1topp2* displayed an ectopic suberization pattern indistinguishable from *topp2* single mutants (Fig. 4g), demonstrating that *TOPP2* is epistatic to *SBG1* and that SBG1 acts upstream of TOPP2 in this regulatory cascade. These findings support a model in which SBG1 inhibits TOPP2 activity, thereby alleviating its negative effect on ABA signaling and promoting suberization of endodermal cells. To further test this hypothesis, we generated plants overexpressing SBG1-mVenus mutated in either the RVxF motif (SBG1_SM) or in both SILK and RVxF motifs (SBG1_DM). In contrast to the WT construct, neither mutant induced suberin deposition (Fig. 4i). Moreover, both mutants showed reduced suberin induction in response to ABA, suggesting a partial dominant-negative effect (Extended Data Fig. 11e). Together with the *in vitro* analysis, these results confirms that SBG1’s positive role in suberization depends on its physical interaction with TOPP2 and inhibition of its phosphatase activity. We therefore identify SBG1 as a key regulator of endodermal suberization acting through inhibition of TOPP phosphatases.

## Discussion

Suberin deposition in roots has attracted growing attention due to its role as a protective barrier against drought and pathogens and as a potential contributor to underground carbon storage. In particular, suberization of the endodermis and exodermis forms a dynamic barrier for water and nutrients^2,8,25,36–40^. Key regulators of endodermal suberin deposition are conserved across vascular plants and were instrumental in the evolution of seed plants^5,38,41,42^. While substantial progress has been made in understanding the molecular pathway of suberin biosynthesis and regulation, most insights come from the *A. thaliana* reference accession Col-0. Moreover, no unbiased, forward genetic approach, aiming at identifying unknown players in suberization had been performed. Previous studies on natural variation in suberin have focused on peridermal suberization in older roots^43^, linking chemical diversity in suberin to soil properties and moisture gradients, identifying 94 putative loci, with verification for *GPAT6*, which has a demonstrated role in suberin biosynthesis^27,43^. Here, we targeted the genetic diversity underlying the highly plastic endodermal suberization to uncover mechanisms involved in nutrient acquisition and stress adaptation. Our screen revealed a strong association between endodermal suberization and accumulation of Rb, Mo, Cu and Zn in shoots grown on agar plates. Unexpectedly, K and Na, classical markers of endodermal suberin levels in Col-0, were not affected, emphasizing that Col-0 may not be representative of species-wide responses, particularly for traits linked to root developmental physiology, as shown before for Zn and P deficiencies responses and for root growth response to hormones^44,45^. While previous studies in Col-0 linked endodermal suberin to B, Rb, Cu and Mo accumulation^3,13,18^, a consistent impact on Zn was not shown before. Accessions with enhanced suberin accumulated less Zn, and several displayed increased sensitivities to Zn limitation, a phenotype that was either unchanged or alleviated when suberin was degraded. These findings align with earlier observations that Zn limitation downregulates suberization, a response proposed to facilitate Zn uptake^2^. Together, these results position endodermal suberin as a contributor to Zn homeostasis, with implications for nutrient adaptation strategies in plants, and highlight an accession-dependent trade-off between protective barrier function and nutrient acquisition.

Our GWAs analysis identified multiple loci associated with endodermal suberization, including two that surpassed the Bonferroni-corrected significance threshold and more than 20 additional loci exceeding the Benjamini–Hochberg threshold. Strikingly, none of the previously known regulators of suberin deposition were found among these loci, underscoring the strength of this unbiased approach in uncovering novel genetic determinants. Focusing on the strongest association on chromosome 1, we identified *SBG1*, a previously unannotated gene whose naturally occurring allelic variation in its 3’UTR impacts the pattern of suberin deposition across accessions, suggesting a major role in suberin-dependent environmental adaptation. Supporting a direct role in suberization, *SBG1* is expressed in the endodermis and is transcriptionally induced by ABA, a well-established hormonal regulator of endodermal suberin deposition, during both stress responses and root development^2,3,8,41^. Consistent with its expression pattern, *SBG1* regulates ABA-induced endodermal suberization and has a broad effect on the transcriptional response to ABA in roots suggesting a regulatory role in the ABA-response. We found that SBG1physically interacts with the protein phosphatases TOPPs, particularly TOPP2, which is constitutively expressed in the endodermis and inhibits its activity. Since TOPPs are known negative regulators of ABA signaling^34,35^, our findings suggest that SBG1 interferes with TOPP activity, thereby locally relieving ABA repression and promoting suberin formation. The enhanced suberin phenotype observed in the *topp2* mutant, which phenocopies WT *SBG1* constitutive expression, and is epistatic to *sbg1* further supports TOPP2 as a central target of SBG1-mediated suberin regulation. Together, our results define a novel regulatory module that fine-tunes ABA signaling in the endodermis to control suberin deposition. Given the evolutionary conservation of SBG1 across angiosperms (Extended Data Fig. 12a,b), this mechanism likely represents a fundamental and broadly relevant strategy for regulating root barrier properties in diverse plant species.

Together, our findings highlight the power of unbiased forward genetic approaches to uncover previously unknown regulators of endodermal suberin deposition. By revealing the genetic basis underlying natural variation in suberin deposition, we identify SBG1 as a central, previously uncharacterized regulator of this key trait. This work not only advances our understanding of the mechanisms governing root barrier formation but also provides a new conceptual and molecular framework to manipulate ABA response and suberin deposition for improving plant stress resilience.

## Methods

### Plant material

The 284 *Arabidopsis* accessions used in this study are listed in Supplementary Table 1. These accessions represent a subset of the 1001 Genomes Project^46^, including 185 of the 195 Salk accessions, 80 Max-Planck Institute accessions, and the 19 founder accessions used to generate the MAGIC (Multiparent Advanced Generation Inter-Cross) population^47^. The ER-GFP marker line was described previously^48^. The T-DNA insertion lines *sbg1-1* (SALK_029983C), *frs6-2* (SALK_114017C)^49^, *at1g52540* (GK-090F10), *at1g52550* (SAIL_681_G01), *pldα2* (GK-212E06), *rbl5* (SALK_079189C), *at1g52590* (SAIL_894_H03.C), *at1g52600* (SALK_062346C), *topp1* (SALK_057537)^34,50^, *topp2* (GK-187C10)^35^, *topp6* (SALK_093747C)^50^, *topp7* (SALK_023073C)^50^, *topp8/aun2-1* (SALK_137888)^51^ and *topp9/aun1-1* (SALK_045433C)^50,51^ were obtained from the Nottingham *Arabidopsis* Stock Centre, and genotyping primers are provided in Supplementary Table 6. Additional mutants (*sbg1-2, sbg1-3, sbg1-4, sbg1;sbg1l1 and quad-sbg1*) were generated via CRISPR-Cas9 in specified accessions using guide RNAs described in the Constructs section. The double mutants *sbg1-1/topp2* and *sbg1-2/topp2* were generated by crossing of corresponding single mutants and homozygous double mutants were isolated by genotyping and sequencing. The following transgenic lines were developed in this study: *UBQ10::SBG1_acc* (*SBG1* from specified accessions), *UBQ10::SBG1-mVenus*, *UBQ10::SBG1_SM-mVenus, UBQ10::SBG1_DM-mVenus, SBG1::NLS-3xmVenus*, *SBG1::SBG1*, *SBG1::SBG1-mVenus, UBQ10::SBG1-mCherry*, *UBQ10::CYP86A1-mScarletI*/*UBQ10::SBG1-mVenus*, *UBQ10::TOPP2-mScarletI* /*UBQ10::SBG1-mVenus* and *TOPP2::NLS-3xmvenus*. The corresponding gene IDs are: *SBG1*, *AT1G52565*; *SBG1L1*, *AT3G15760*; *SBG1L2*, *AT2G35850*; *SBG1L3*, *AT3G52360*; *TOPP1*, *AT2G29400*; *TOPP2*, *AT5G59160*; *TOPP6*, *AT5G43380*; *TOPP7*, *AT4G11240*; *TOPP8*, *AT5G27840*; *TOPP9*, *AT3G05580*; *GPAT5*, *AT3G11430*; *CYP86A1*, *AT5G58860*; *MYB41, AT4G28110*; *FRS6, AT1G52520*; *PLDα2, AT1G52570*; *RBL5, AT1G52580*; *I2, AT5G52200*.

### Constructs and plant transformation

CRISPR-Cas9 constructs targeting *SBG1* were designed using Benchling (https://www.benchling.com/) (Supplementary Table 6). Oligonucleotides for guide RNAs were annealed, digested by BbsI, and ligated into the pEN-Chimera vector^52^. The resulting entry constructs were recombined using the Gateway Technology® (Thermo Fisher Scientific) into a destination vector containing the *UBQ10::SpCas9* and a FAST Red selection marker^53^. Entry clones for expression constructs were generated using Gibson assembly following PCR amplification. PCR products were cloned into a BamHI-digested pUC57-L4_EcoRV-XbaI-BamHI_R1 or into a NotI-digested pDONR221-*L1_SpeI-BglII-NotI-SphI-EcoRI_L2.* Newly generated entry clones are: *L4-proSBG1-R1* (3000bp promote*r*), *L4-proTOPP2-R1* (1322bp), *L1-SBG1-L2* (genomic *SBG1* without stop codon from Col-0), *L1-SBG1_acc-L2* (genomic *SBG1* including 1500 bp of 3’ region from specified accession), *L1-TOPP2-L2* (genomic *SBG1* without stop codon), *L1-CYP86A1-L2* (CDS). *L1-SBG1_SM-L2* and *L1-SBG1_DM-L2*_were synthesized and then recombined in the same way as for *UBQ10::SBG1-mVenus*. For yeast two-hybrid assay, CDS of *SBG1* 88-390bp, *TOPP1* and *TOPP2* were cloned in pDONR221 by Gateway BP reaction. *L1-SBG1_SM (V63A/F65A)-L2* and *L1-SBG1_DM (V63A/F65A, C46A/L47A)-L2* 88-390bp were synthesized. Then they were recombined into pGADT7 or pGBKT7 (TaKaRa) by Gateway LR reaction. For protein expression in *Escherichia coli* (*E. coli*), the CDS of *SBG1_Col-0*, *SBG1_Tsu-0, SBG1_DM* and *TOPP2* were cloned by Gibson assembly in pETM41 vector harboring an N-terminal 6xHis-MBP tag. Previously described entry clones include *L4-pUBQ10-R1*^54^ (promoter), *L1-nls-3xmVenus-L2*^55^*, R2-mCherry-L3*^55^, *R2-mVenus-L3*^56^ and *R2-mScarletI-L3*^57^ (CDS), and *R2-tHSP18.2-L3*^58^ and *R2-tNOS-L3*^59^ (terminators). Expression clones were generated using the Multisite Gateway Technology (Life Technologies, Carlsbad, CA, USA) by recombination of entry clones into the destination vectors pFR7m34GW (FAST Red) or pFG7m34GW (FAST Green)^58^. Primers used for cloning are listed in Supplementary Table 5. Final constructs were introduced into *Arabidopsis* using *Agrobacterium tumefaciens* strain GV3101 and the floral dip method^60^. For each construct, at least 14 T1 seeds were selected based on fluorescence (FAST Red or FAST Green)^61^. Segregation ratios, gene expression levels, and phenotypes were analyzed across 14 independent T2 lines. All experiments were conducted on T3 homozygous lines with mono-insertional events unless otherwise noted.

### Growth conditions

For *in vitro* experiments, seeds were surface sterilized using chlorine gas and sown on square plates. Unless stated otherwise, seedlings were grown on half-strength MS (Murashige and Skoog) medium (pH 5.7) containing 0.8 % agar without sucrose. For nutrient deficiency assays, the medium was prepared using a 20x macronutrient stock solution (Sigma-Aldrich, M0654), 2000x individual micronutrient stock solutions consisting to reach a final concentration of 9.4 mM KNO*3*, 10.3 mM NH₄NO₃, 625 mM KH*2*PO*4*, 1.49 mM CaCl*2*, 0.75 mM MgSO*4*, 50 µM H*3*BO*3*, 0.05 µM CuSO*4*·5 H*2*O, 50 µM MnSO*4*·H*2*O, 0.5 µM Na*2*MoO*4*·2 H*2*O, 15 µM ZnSO*4*·7 H*2*O, 2.5 µM KI, 0.05 µM CoCl*2*·6 H*2*O, 50 µM Fe-EDTA. MES (0.5 g/L) was added and pH adjusted to 5.7 with KOH. For nutrient-deficient conditions, the specified nutrient was omitted from the medium. For the suberin screening in 284 accessions, seeds were stratified for 7 days at 4°C to minimize germination variability. For other experiments, stratification was performed for 2 to 3 days at 4°C. Seedlings were grown vertically in growth cabinets set to 22°C under continuous light (∼100 µE) and 40% relative humidity. For propagation and salt treatment in soil, plants were grown under long-day conditions (16 h light/8 h dark) at 20 ± 2°C, with a light intensity of 150-180 µE and 60% relative humidity. Experiments were conducted on 5-day-old seedlings unless otherwise specified.

### Suberin staining and pattern quantification

For suberin histological analysis, seedlings were whole-mount stained as described previously^3^. Briefly, whole seedlings were immersed in Fluorol Yellow 088 (0.01% w/v, lactic acid; Chem Cruz, Cat# sc-215052) and incubated at 70°C for 30 minutes in the dark. After staining, seedlings were washed twice with water and counterstained with aniline blue (0.5% w/v; Sigma-Aldrich, Cat# 415049) for 30 minutes at room temperature in the dark, followed by two additional water washes. For imaging, seedlings were mounted on slides in 50% glycerol and scanned using a ZEISS Axio Zoom.V16 stereomicroscope equipped with a GFP filter (excitation: 450-490 nm, emission: 500-550 nm). Full root imaging at high resolution was achieved using a tile-scanning approach. The region of interest (ROI) was first defined during a low-resolution scan (7x magnification). Focus points were manually selected along the root to ensure consistent focus, and high -resolution images (63x magnification) were captured as overlapping tiles (10% overlap) that were subsequently stitched together. Suberin pattern quantification was performed by measuring the lengths of three distinct suberin zones along the root (non-suberized, patchy, and fully suberized) using Fiji^62^ software as previously described^3^.

### Correlations of suberin pattern with climate factors

Climate data for the *Arabidopsis* accessions analyzed in this study were retrieved from AraCLIM (https://github.com/CLIMtools/AraCLIM)^63^. Relevant climate variables were selected and extracted for the geographic origin of the characterized accessions. The genetic similarity between accessions was taken into account in the correlation, to avoid any confounding effect of population structure in the relationship between the endodermal suberization and the climate factors. To correct for the population structure, genotyping information for 261 of the 284 *A. thaliana* accessions in this study was extracted from the 1001 genomes project^46^ and from the imputed RegMap data^64^. The variant call format (VCF) from the two sources were merged using vcftools v1.19^65^ and the final VCF filtered for biallelic sites, a minor allele frequency of 0.05 and a maximum of 10% missing data per site with vcftools v0.1.16^66^. This genotyping matrix was summarised by PCA using the SNPRelate v1.42.0 R package^67^, and the two first principal components PC1 (accounting for 17.45% of variance) and PC2 (accounting for 7.8% of variance) were taken as covariates in the linear model (lm) assessing the link between suberin data and environmental variables: suberin∼ environmental variable + PC1 + PC2. This linear model was fitted on raw, as well as on scaled environmental data, to enable the comparison of the slopes of the lm between different environmental variables. The PC1 and PC2 covariates representing population structure were never significantly correlated with the suberin data. Environmental variables were considered to be significantly associated with suberin when the *P*-value for their regression coefficient was below 0.05, based on a two-sided t-test. The partial R² for the environmental variable presented in Supplementary Table 2 was calculated by subtracting the R² of the linear model (with environmental variable, PC1 and PC2 as predictors) by the R² of a linear model only taking PC1 and PC2 as predictors. These analyses were performed on R v4.5.0.

### Propidium iodide staining and uptake assay

Propidium iodide (PI) was used to counterstain cell walls and as an apoplastic tracer for evaluating Casparian strips functionality, as described previously^24^. Seedlings were live stained with 15 µM PI in darkness for 10 minutes, followed by two washes with water. For the PI uptake assay, the functionality of the apoplastic barrier was assessed by counting the number of endodermal cells beyond the onset of elongation where PI uptake was blocked by Casparian strips. The onset of elongation was defined as the point where the first endodermal cell exhibited a length at least three times its width. Observations were made using a Leica DM6 B epifluorescence microscope equipped with an I3 filter at 20x magnification.

### Confocal microscopy and image analysis

Confocal microscopy imaging was performed using Leica SP8, Zeiss LSM 780, and Zeiss LSM 800 microscopes. Excitation and detection wavelengths were set as follows: for Leica SP8 system, GFP/mVenus (488 nm, 500-546 nm), PI/mCherry (552 nm, 585-617 nm); for Zeiss LSM 800, GFP/mVenus (488 nm, 500-546 nm), PI (561 nm, 585-617 nm); for Zeiss LSM 780 mVenus (514 nm, 515-561 nm), PI (514 nm, 585-679 nm). Scanner and detector settings were maintained constantly across all experiments to ensure consistency. Image analyzed was conducted using ZEN2.3 software (ZEISS) and Fiji^62^ (http://fiji.sc/Fiji).

### Ionomic analysis

Elemental content in 7-day-old leaves was measured using Inductively Coupled Plasma Mass Spectrometry (ICP-MS). Dried leaves were weighed and transferred to 15 mL metal-free Falcon tubes (VMR). Samples were digested with 300 µL of 67% nitric acid (Fisher Chemicals) containing 100 ppb indium as an internal standard and incubated in a dry block heater at 100°°C for 2 hours. After cooling, digests were diluted to 15 mL with Milli-Q water (Merck Millipore). Elemental analysis was performed on an Agilent 7700X ICP-MS with an autosampler operating in helium collision mode. Scandium (2 ppb) was added as an on-site internal standard. Fifteen elements (B, Na, Mg, P, S, K, Ca, Mn, Fe, Co, Cu, Zn, Rb, Sr and Mo) were monitored. Calibration standards were prepared using single-element standards solutions (Inorganic Ventures; Suisse Technology Partners Ltd.). A standard reference material (NIST1515, apple leaves, Sigma-Aldrich) was included to validate analytical methods. Element concentrations in the samples were calculated using the external calibration method, normalized to the dry weight, and processed with the instrument software.

### Chlorophyll quantification

Shoot materials were collected in 1.5 mL Eppendorf tubes after measuring their fresh weight. Then dimethylformamide (DMF) with (volume:weight = 10:1) was added to the samples followed by centrifugation with 15,000 rpm at room temperature for 1 min. The samples were stored at 4℃ overnight. After 3-min centrifugation with 15,000 rpm at room temperature, the absorbance at 647 nm and 664 nm were measured using a Tecan Spark plate reader. The total chlorophyll content was calculated based on Porra’s method^68^.

### Genome-wide association analysis

The GWAS was performed on GWA-Portal (https://gwas.gmi.oeaw.ac.at/) with the data of the fully suberized zone quantified in 265 accessions (Supplementary Table 1) and Imputed Full sequence Dataset corresponding of a combination of the 250K SNP dataset and the 1001 genomes dataset using imputation (TAIR9)^69^. An accelerated mixed model (AMM) was applied for the GWAS and a significance threshold of 10% was used after Bonferroni-Hochberg correction for multiple tests. Gene candidates in the 15 kb distance to the top SNPs with -log_10_*P* > 6 and minor allele count (mac) ≥ 15 are listed in Supplementary Table 1.

### RNA secondary structure prediction

The mRNA secondary structure prediction for the 3’UTR of *SBG1* from different accessions was performed on RNAfold web server (http://rna.tbi.univie.ac.at/cgi-bin/RNAWebSuite/RNAfold.cgi). The prediction followed the default settings with the fold algorithms of minimum free energy (MFE) and partition function and avoiding isolated base pairs, dangling energies on both sides of a helix and RNA energy parameters from Turner model^70^.

### Suberin monomer analysis

Suberin monomers analysis in roots was conducted by gas chromatography-mass spectrometry (GC-MS) following a previously described protocol^71^. Roots from 5-day-old seedlings (200-300 per replicate) were harvested and incubated in isopropanol containing 0.01% butylated hydroxytoluene (BHT) at 85°C for 10 minutes. Delipidization was performed twice (16 hours and 8 hours) using extractions with chloroform-methanol (2:1), chloroform-methanol (1:1), and methanol with 0.01% BHT, with samples continuously agitated. The delipidized samples were vacuum-dried for 3 days before depolymerization via base catalysis. For depolymerization, dried plant material was subjected to trans-esterified in a reaction medium (20 mL) composed of 3 mL methyl acetate, 5 mL of 25% sodium methoxide in dry methanol, and 12 mL dry methanol. Methyl heptadecanoate (5 mg) and w-pentadecalactone (10 mg) were added as internal standards per sample. The samples were incubated at 60°C for 2 hours, followed by the addition of 3.5 mL dichloromethane, 0.7 mL glacial acetic acid, and 1 mL 0.9% NaCl in 100 mM Tris-HCl buffer (pH 8.0). Samples were vortexed for 20 seconds and centrifuged at 1500 x g for 2 minutes. The organic phase was collected, washed with 2 mL 0.9% NaCl, dried over sodium sulfate, and concentrated under a nitrogen stream. The suberin monomers were derivatized with N,O-bis(trimethylsilyl)trifluoroacetamide (BFTSA) in pyridine (1:1, v/v) at 70°C for 1 hour before GC-MS analysis. Samples were injected in hexane onto an HP-5MS column (J&W Scientific) using an Agilent 6890N GC Network systems coupled to a mass spectrometer and flame ionization detector. The oven temperature was programmed as follows: 50°C for 2 minutes, an increase of 20°C/min to 160°C, 2 °C/min to 250°C, and 10°C/min to 310°C, held for 15 minutes. The amounts of unsubstituted C16 and C18 fatty acids were excluded from the analysis due to their widespread occurrence in plants and the environment.

### Protein sequence alignment and phylogenetic analysis

The protein sequences of SBG1, SBG1L1, SBGL2 and SBG1L3 were aligned with Clustal Omega methods on SnapGene (https://www.snapgene.com/). The protein sequences of SBG1 orthologs were acquired by NCBI’s protein blast to select the protein with the highest similarity across different species. Sequences were subsequently aligned with the Clustal Omega methods in (https://www.ebi.ac.uk/jdispatcher/msa/clustalo)^72^. The phylogenetic analysis was conducted on ClustalW2 with Phylip output format and Neighbour-joining clustering methods (https://www.ebi.ac.uk/jdispatcher/phylogeny/simple_phylogeny)^72^.

### qRT-PCR

Gene expression analyses were performed using roots from 7-day-old seedlings, with approximately 60 roots pooled per biological replicate (3 biological replicates per genotype and conditions). After harvesting, roots were immediately frozen in liquid nitrogen and subsequently ground to a fine powder using a tissue lyser. Total RNA was extracted using the TRIzol reagent (Ambion) and further purified with the RNeasy MinElute Cleanup Kit (Qiagen). Reverse transcription was performed with the Maxima First Strand cDNA Synthesis Kit (Thermo Scientific). Quantitative real-time PCR (qRT-PCR) was conducted on an Applied Biosystems QuantStudio5 thermocycler using SYBR Green master mix (Applied Biosystems). The expression levels were normalized to the housekeeping gene *Clathrin AP4M* (*AT4G24550*), and relative expression was calculated using the 2^-ΔΔCt^ method^73^. Primers used for qRT-PCR are detailed in Supplementary Table 6.

### RNA sequencing and analysis

Roots of 6-day-old seedlings were harvested and frozen by liquid nitrogen. The RNA extraction and purification were performed following the same protocol as for qRT-PCR. RNA samples were sent to Novogene for sequencing and analysis based on Illumina platform with Pair-end 150 bp strategy and 6 G data acquirement. After quality controls, high-quality reads were mapped to *Arabidopsis thaliana* TAIR10 genome with HISAT2 and the matrix of counts per gene was produced. The differential expression analysis was performed with DESeq2 and subjected for a multiple testing Benjamini and Hochberg correction with 5% FDR. The differential expressed genes (DEGs) are listed in Supplementary Table 4. For the GO enrichment analysis, all the genes with *P*_adj_ < 0.05 was selected and analysed on g:Profiler (https://biit.cs.ut.ee/gprofiler/gost).

### IP-MS analysis

Roots of 7-day-old seedlings expressing SBG1-mVenus and ER-GFP were harvested for protein extraction. After grinding the total protein was extracted with 50 mM Tris-HCl pH 7.5, 150 mM NaCl, 2 mM EDTA, 5 mM Dithiothreitol, 1 mM PMSF, 1% Triton X-100 and protease inhibitor cocktail (Roche). To immunoprecipitate mVenus- or GFP-tagged proteins, GFP-Trap Magnetic Agarose beads (Chromotek) were added to the supernatant of protein extracts. Beads were first washed 3 times with 50 mM Tris-HCl pH 7.5, 150 mM NaCl, 2 mM EDTA, 0.01% Triton X-100, then 3 more times with 50 mM Tris-HCl pH 7.5, 150 mM NaCl. The beads were kept in the second washing buffer. To prepare samples for proteomic analysis, the remaining buffer was removed, and the beads were suspended in 100 µL 1 M Urea in 50 mM Ammonium bicarbonate. Samples were reduced using a final concentration of 10 mM DTT at RT for 30 min and alkylated with 20 mM iodoacetamide at RT for 30 min in the dark. Alkylation reactions were quenched with a half amount of DTT used for the reduction and incubated 10 min at RT in the dark. Finally, beads were incubated with 300 ng Trypsin (Promega, USA) in 1 mM hydrochloric acid and 25 mM ammonium bicarbonate and digested overnight at 25°C. Afterward, digestion was stopped by adding 10% TFA to reach a final concentration of 0.5%, centrifuged and deposited on a magnetic rack to transfer the supernatant into a fresh tube. Tryptic peptides were desalted using MonoSpin C18 columns (GL Sciences Inc., Japan) and eluted from C18 columns using 0.1% TFA in 50% acetonitrile and dried in a centrifugal vacuum concentrator. Tryptic peptides were dissolved in 0.1% (v/v) formic acid in 2% (v/v) acetonitrile for mass spectrometry (MS) analysis.

For MS analysis, samples were measured on an Easy Nano LC Orbitrap Fusion System equipped with a nanospray flex™ ion source (Thermo Fisher Scientific, USA). Peptides were separated on a 1.9 μm particle, 75 μm inner diameter, 15 to 20 cm filling length homemade C18 column. A flow rate of 300 nL/min was used with a 2 h gradient 3–30% solvent B. The gradient followed with two rounds of washing steps, in each step, the gradient switched to 90% solvent B in 1 min and kept for another 5 min, and switched to 3% solvent B in 1 min and kept for another 5 min. In the second round of washing the 90% B solvent was kept for 2 min, an extra 14 min of 3% solvent B was kept for system equilibration. Solvent A was 0.1% (v/v) formic acid in LC/MS grade water and solvent B was 0.1% (v/v) formic acid in 100% (v/v) acetonitrile. The ion source settings from Tune were used for the mass spectrometer ion source properties.

For data-independent acquisition (DIA), data were acquired with 1 full MS and 41 overlapping isolation windows constructed covering the precursor mass range of 350–1400 m/z. For full MS, Orbitrap resolution was set to 120,000. Automatic Gain Control (AGC) target was set to custom with normalized AGC target set to 50%, and maximum injection time (IT) was set to 100 ms. DIA segments were acquired at a resolution of 30,000. AGC target was set to custom with normalized AGC target set to 200%, and a 54 ms maximum IT. HCD fragmentation was set to normalized collision energy of 28%. For protein identification and quantification, Raw files from DIA measurements were analyzed using the directDIA workflow in Spectronaut software (Biognosys, Switzerland) with default settings. Identification was performed using the Uniprot reference proteome UP000006548 for *Arabidopsis thaliana*.

### Co-immunoprecipitation and immunoblot analysis

For co-immunoprecipitation (Co-IP) assays, 7-day-old transgenic seedlings co-expressing SBG1-mVenus and CYP86A1-mScarletI or TOPP2-mScarletI were harvested and ground in liquid nitrogen. Total proteins were extracted with 50 mM Tris-HCl pH 7.5, 150 mM NaCl, 2 mM EDTA, 5 mM Dithiothreitol, 1 mM PMSF, 1% Triton X-100 and protease inhibitor cocktail (Roche). After four centrifugations at 16,000 × g for 15 min at 4 °C, 100 µL of the extracts was kept as input for the immunoblot. For immunoprecipitation the remaining supernatant was incubated for 2 h at 4°C with GFP-Trap Magnetic Agarose beads (Chromotek) with gentle rotation. Beads were then washed 3 times with 50 mM Tris-HCl pH 7.5, 150 mM NaCl, 2 mM EDTA, 0.01% Triton X-100, then 3 more times with 50 mM Tris-HCl pH 7.5, 150 mM NaCl and eluted in 2x Laemmli buffer (100 µL buffer into 50 µL beads) by boiling for 5 minutes at 95℃. Input (25 µL, boiled in 2x Laemmli 5 minutes at 95 °C) and immunoprecipitated fractions (2 µL) were separated by SDS-PAGE and transferred to PVDF membranes. Immunoblots were probed with anti-GFP antibody (1:4000, JL-8, Clontech) or anti-RFP antibody (1:4000, 6G6, Chromotek) to detect mVenus and mScarletI respectively, followed by anti-mouse IgG (H+L) HRP-conjugated secondary antibodies (1:10000, W4021, Promega) and detection using SuperSignal West Femto Maximum Sensitivity Substrate (34095, Thermo Scientific). Signals were visualized with a ChemiDoc imaging system (Amersham Imager 680).

### Alphafold3 analysis

Protein interaction between SBG1, SBG1L1, SBG1L2, SG1L3, I2 and TOPP2 was predicted by AlphaFold3 model (https://alphafoldserver.com/). Full protein sequences of SBG1s, I2, TOPP2 and 2x Mn^2+^ were entered as input for the prediction. pLDDT coded structure and PAE plots indicating the quality of the prediction are shown in Extended Data Fig. 7 d,e. Final models were imported and modified with PyMOL V3.1.0 (https://www.pymol.org/).

### Yeast two-hybrid

Yeast two-hybrid analyses were conducted using Matchmaker Gold Yeast Two-Hybrid System (Clontech; TaKaRa, Tokyo, Japan) according to the manufacturer’s protocols. Yeast strain (Y2HGold) was co-transformed with the respective AD and BD plasmids and transformants were selected on SD-LW (leucine, tryptophane). For interaction analysis, 5µl of yeast culture dilutions in the indicated concentration range were spotted on both SD-LW and SD-LWH (leucine, tryptophane, histidine) plates and incubated at 28°C for 3 d.

### Protein expression and purification

The protein expression constructs of SBG1_Col-0, SBG1_Tsu-0, SBG1_DM and TOPP2 with N-terminal 6xHis-MBP were first transformed into *E. coli* BL21 (DE3) strain. Then, two liters of LB media with kanamycin were inoculated with 3 mL overnight cultures for around 4 h at 37 ℃ until the OD600 reached 0.6. To induce protein expression, 0.5 mM isopropyl β-D-1-thiogalactopyranoside (IPTG) was added and the culture was incubated at 18 ℃ overnight. The cells were harvested by centrifugation and stored at −80 ℃. For protein purification, the harvested cells expressing SBG1_Col-0, SBG1_Tsu-0, SBG1_DM or TOPP2 were resuspended in a lysis buffer containing 50 mM HEPES pH 8.0, 500 mM NaCl, 1 mM MnCl_2_, 2 mM DTT and 5% glycerol. After cell lysis and centrifugation, the supernatant was filtered and applied to a 5 ml Hitrap MBP column equilibrated in lysis buffer. Proteins were eluted using a buffer containing 50 mM HEPES pH 8.0, 500 mM NaCl, 20 mM maltose, 1 mM MnCl_2_ and 2 mM DTT. Fractions containing SBG1 or TOPP2 were pooled, concentrated to 0.5 ml and loaded on a Superose 6 Increase 10/300 GL gel filtration column (GE Healthcare Life Sciences) pre-equilibrated in 25 mM HEPES pH 8.0, 500 mM NaCl, 1 mM MnCl_2_ and 2 mM DTT. Monodisperse peak of the target protein was collected, flash-frozen in liquid nitrogen and stored at −80 ℃.

### Analytic gel filtration

To detect protein complex formation, 100 µg of purified TOPP2 was mixed with either 100 µg of wild-type or mutated SBG1 in a buffer containing 25 mM HEPES pH 8.0, 500 mM NaCl, 1 mM MnCl_2_ and 2 mM DTT. The protein mixture was incubated at 4 °C for 40 min followed by gel filtration on a Superose 6 3.2/300 column (GE Healthcare Life Sciences) in the same buffer. Fractions of eluted peaks were collected and analyzed by SDS–PAGE with Coomassie staining.

### Phosphatase activity assay

The phosphatase activity of TOPP2 with SBG1 was measured as previously described^30^, using the phosphorylated peptide H3pT3 (Histone 3 pT3: NH2-AR(pThr)KQTARKSTGGKAPRKQL-COOH, GenScript) as a substrate. Malachite Green reagent (MAK308, Sigma) was used to detect free Pi. Briefly, 25 µL of serially diluted SBG1 protein was mixed with 15 µL of 3.5 nM TOPP2 or reaction buffer (50 mM HEPES pH 8.0, 150 mM NaCl, 1 mM MnCl_2_ and 1 mM DTT; low-signal control) in 96-well plate and incubated at room temperature for 30 min. Then, 10 µL of 500 µM H3pT3 peptide was added into SBG1-TOPP2 mixtures, followed by a 20-minute reaction at 30 ℃. The reaction was terminated by addition of 50 µL Malachite Green reagent and after 30 min the absorbance at 620 nm was measured using a Tecan Spark plate reader. Relative phosphatase activity was calculated by subtracting the background signal (low-signal control) and then normalizing resulting values to TOPP2 control where SBG1 was absent. IC50 curves were generated by fitting the data to a 4-parameter logistic model (Prism V10.6.1).

### Statistical analysis

Statistical analysis and graph generation were performed using GraphPad Prism 10 (https://www.graphpad.com/) except for the correlations with climate variable where the R^74^ v4.5.0 environment was used. All experiments were conducted independently at least 3 times, yielding consistent results, except for the proteomic analysis (2 independent experiments with 3 biological replicates) and the RNA-seq (performed once with 3 independent pools of 250 roots). Nonparametric Spearman correlations were performed on Prism with 95% confidence interval and two-tailed test. Cohen’s *d* was calculated as the difference between the means of the two groups divided by their pooled standard deviation: Cohen’s *d* = (*M*_2_ - *M*_1_) ⁄ *SD*_pooled_, where, *SD*_pooled_ = √(((n_1_-1)\**SD*_1_^2^ +(n_2_-1) \**SD*_2_^2^) ⁄ (n_1_+n_2_-2)). All the datasets were first tested for normal distribution and equality of variances. Depending on these results, comparisons of means were performed using either parametric or non-parametric tests. The tests used are specified in the legends. The PCA and clustering analysis of ionomic profiles were performed on MVApp^75^ (https://mvapp.kaust.edu.sa/).

## Acknowledgments

We are grateful to Robertas Ursache, Lothar Kalmbach and Satoshi Fujita for sharing plasmids. We thank Sylvain Loubéry and the Photonic Bioimaging Center at the University of Geneva for their invaluable assistance with microscopy and image analysis, as well as Sophie Michalet and the ICP-MS center at University of Geneva for support with ionomic analyses. We thank Houming Chen and Zhoubo Hu for their support on IP-MS analysis and AlphaFold predictions and Rodrigo Reis for his advice on RNA structure predictions. Ilona Anton is thanked for her assistance on the physiological analysis of *CDEF1* lines. We thank Richard Chappuis and Roman Ulm for their guidance with yeast two-hybrid. Special thanks go to Laura Peréz Martín, Fabienne Cléard-Karch, Léa Jacquier, and Kevin Robe for their assistance with sample collection for chemical analyses of suberin. We thank Ines Hadj Bachir for bridging the collaboration of bioinformatic analysis. Isabelle Fleury is warmly thanked for her dedicated support in plant growth and propagation. This work was supported by the Sandoz Family Monique De Meuron philanthropic foundation’s program for academic promotion and the Swiss National Science Foundation (PCEGP3_187007) awardee to MB, as well as by the state of Geneva.

