## Extended Figure 1 for "GWAs reveals SUBER GENE1-mediated suberization via Type One Phosphatases"

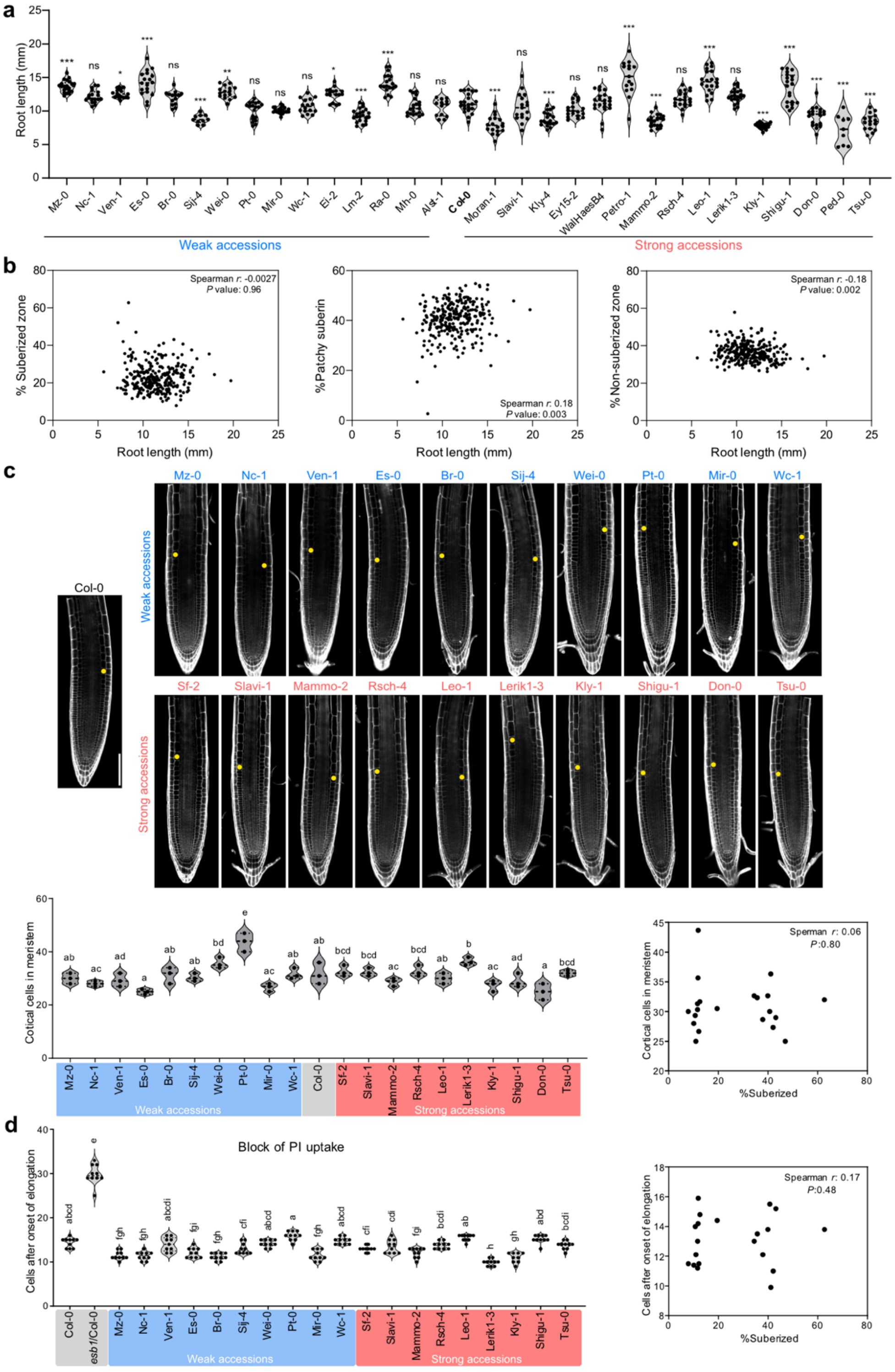


**Extended Data Fig. 1 | Root morphology of extreme accessions.** **a**, Primary root length of 4-day-old Col-0 and 30 extreme accessions selected from Fig.1a. Data are presented as violin plots with individual data points overlaid, $n$ = 10. Thick dashed lines in the violin plots indicate the median, while dotted lines represent the first and third quartiles. Statistical differences were assessed by one-way ANOVA followed by Dunnett's test and are presented for the comparison to Col-0 (*$P$ < 0.05, **$P$ < 0.01, ***$P$ < 0.001). **b**, Correlation analysis between primary root length and the extent of fully suberized, patchy suberized, or non-suberized zones across the 284 accessions. Each dot represents the mean value for one accession. **c**, Root morphology in 5-day-old extreme accessions (upper panels). Cell walls were stained with PI (grey). Yellow dots mark the onset of cortex cell elongation. Scale bar, 100 µm. Violin plots presented as in (**a**) showing the number of cortical cells under division in the meristem, $n$ = 3 (lower, left panel). Statistical differences were assessed by one-way ANOVA followed by Tukey's test; different letters indicate significant differences between genotypes ($P$ < *0.05*). No correlation was found between meristem size and fully suberized pattern ($P$ > 0.05) (lower, right panel). Each dot in the scatter plot represents the mean value for one accession. **d**, Quantification of apoplastic barrier formation in extreme accessions. Violin plots presented as in (**a**) showing the number of cells between the onset of elongation and the site of PI block, $n$ = 10 (left panel). Statistical analyses were performed as in (**c**). No significant correlation was found between the PI blocking and fully suberized pattern ($P$ > 0.05) (right panel). Each dot in the scatter plot represents the mean value for one accession.
