## Extended Figure 2 for "GWAs reveals SUBER GENE1-mediated suberization via Type One Phosphatases"

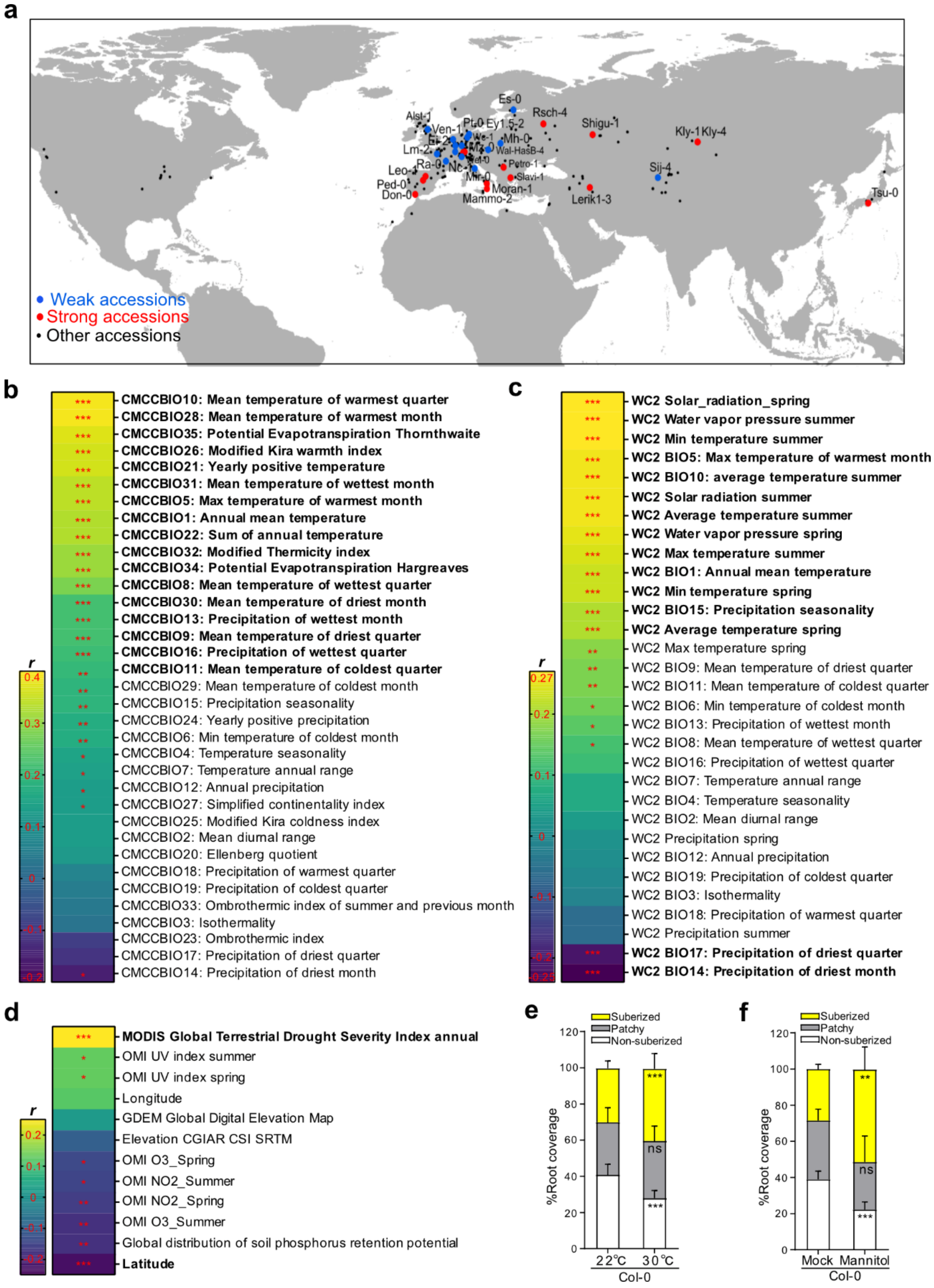


**Extended Data Fig. 2 | Suberin variations correlate with climate variables.** **a**, Geographical origin of the 284 accessions. Weak and strong accessions are highlighted in blue and red, respectively. **b-d**, Correlations between the extent of the suberized zone in 284 accessions and climate variables obtained from 3 different datasets: CMCC (**b**), WorldClimate (**c**) and OMI (**d**). Climate variables with an absolute correlation coefficient (|r|) ≥ 0.2 are shown in bold. A two-side Student's *t*-test was used to assess significance (**P* < 0.05, ***P* < 0.01, ****P* < 0.001). Data for (**b-d**) is provided in Supplementary Table 2. **e**, **f**, Suberin deposition pattern in Col-0 with or without heat (**e**) or osmotic (**f**) stresses. For the treatments, seedlings were grown on 1/2 MS at 22 °C for 4 days, then they were transferred to either 30 °C or 1/2 MS supplemented with 200 mM mannitol for 24 h. Data are presented as stacked columns showing the mean percentage of each suberization zone + s.d., *n* = 10. Student's t-test was used to assess significance for the suberization zones (***P* < 0.01, ****P* < 0.001, ns, not significant).
