## Extended Figure 3 for "GWAs reveals SUBER GENE1-mediated suberization via Type One Phosphatases"

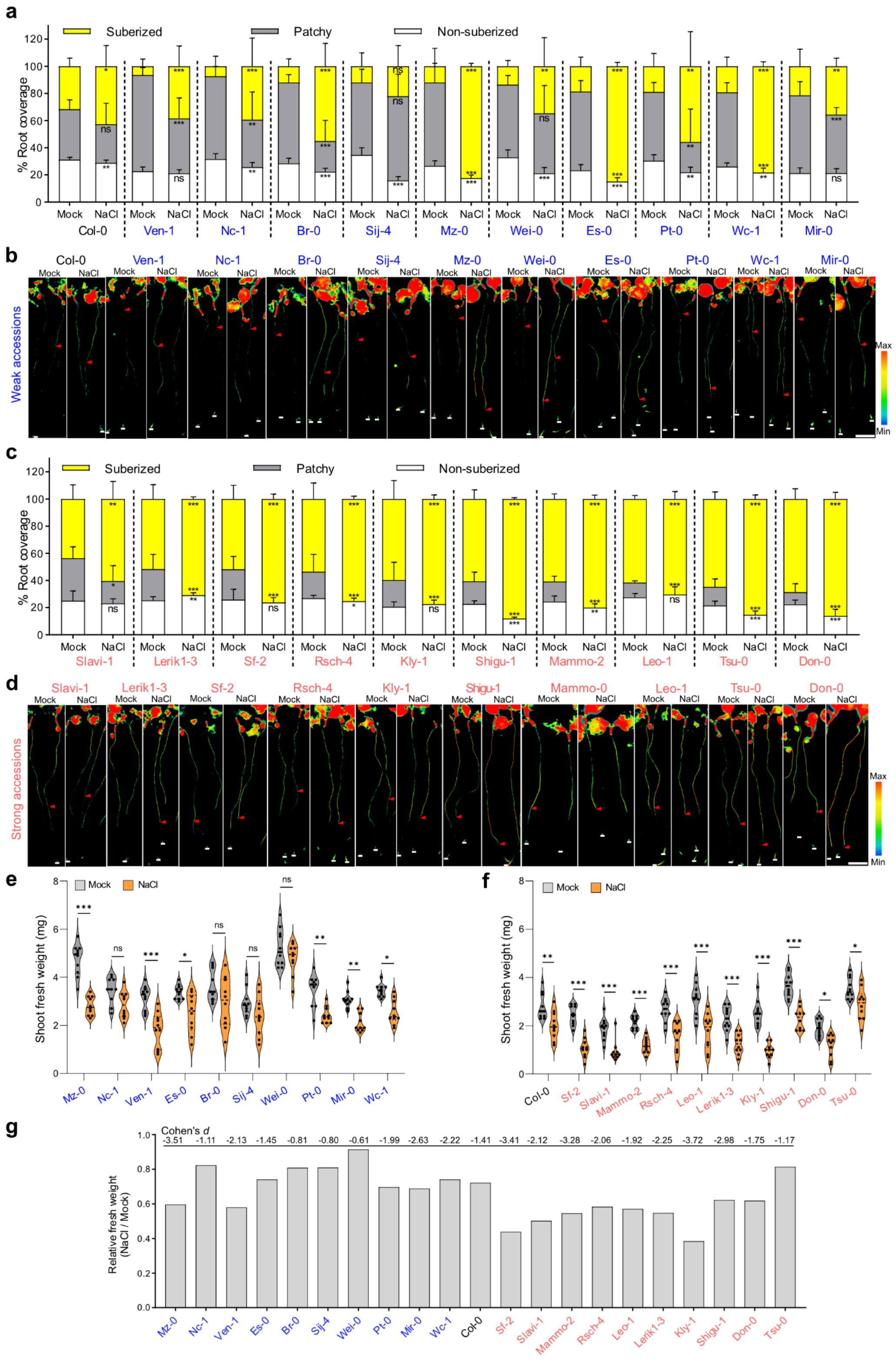

**Extended Data Fig. 3 | Effect of salt stress on extreme accessions. a**,**c**, Suberin quantification in Col-0 and 10 weak (**a**) or 10 strong (**c**) accessions under control (mock) or moderate salt stress (85 mM NaCl). 4-day-old seedlings were transferred to mock or NaCl-containing plates for 22 h prior to suberin staining. Data are presented as stacked columns showing the average percentage of each suberization zone + s.d., with $n$ = 10. Accessions are sorted by percentage of suberized zone under mock condition. Significance between mock and NaCl conditions for the three suberization patterns was tested using a two-tailed Student's t-test (*$P$ < 0.05, **$P$ < 0.01, ***$P$ < 0.001, ns, not significant). **b**,**d**, Whole-mount images of Fluorol Yellow-stained roots from selected extreme accessions under mock and NaCl conditions from (**a**,**c**). Red arrows indicate the onset of the suberized zone; white lines indicate the root tip. Scale bar, 2 mm. **e**,**f**, Shoot fresh weight of weak (**e**) and strong (**f**) accessions under mock or NaCl conditions. 4-day-old seedlings were transferred to either ½ MS plate or ½ MS plate supplemented with 85 mM NaCl for 4 days. Data are shown as violin plots with individual values overlaid, $n$ = 10. Thick dashed lines in the violin plots indicate the median, while dotted lines represent the first and third quartiles. Two-way ANOVA followed by Sidak's test was used to assess the significant differences between mock and NaCl conditions for each accession (*$P$ < 0.05, **$P$ < 0.01, ***$P$ < 0.001, ns, not significant). **g**, Relative fresh weight of the accessions shown in (**e**,**f**) under mock and NaCl conditions. Cohen's $d$ values are shown on the top of histogram to evaluate the effect sizes by NaCl for each accession.
