## Extended Figure 4 for "GWAs reveals SUBER GENE1-mediated suberization via Type One Phosphatases"

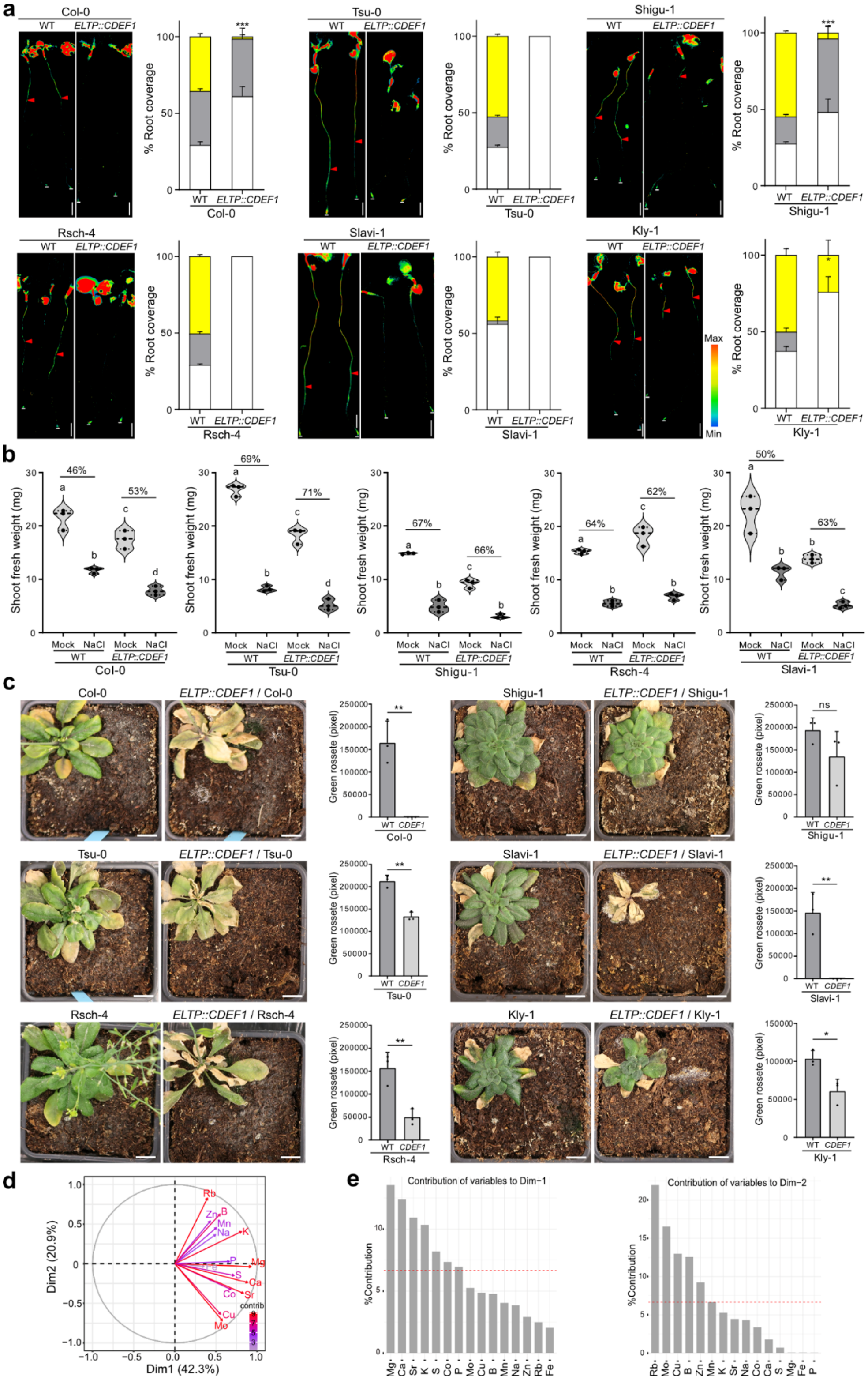


**Extended Data Fig. 4 | Effect of salt stress on *CDEF1* lines in strong accessions backgrounds.** **a**, Whole-mount images of Fluorol Yellow-stained roots and suberin deposition pattern in strong accessions without (WT) or transformed with *ELTP::CDEF1*. Red arrows indicate the onset of suberized zone; white lines mark the root tip. Scale bar, 1 mm. Data are presented as stacked columns showing the mean percentage of each suberization zone + s.d., $n = 10$. Student's test was used to assess significant differences for the suberized zone (***$P < 0.001$, *$P < 0.05$). **b**, Shoot fresh weight of seedlings grown on ½ MS plate for 4 days and then transferred to either ½ MS plates or ½ MS plates supplemented with 85 mM NaCl for 10 days. Data are shown as violin plots with individual values overlaid, $n = 3$ with each individual value corresponding to the average value for 10 shoots. Thick dashed lines in the violin plots indicate the median, while dotted lines represent the first and third quartiles. One-way ANOVA followed by a Tukey's test was used to assess significance; different letters indicate significant differences between conditions or genotypes ($P < 0.05$). The reduction ratios of fresh weight by NaCl treatment are presented on the graphs. **c**, Rosette phenotype of plants grown 17 days and then watered with tap water supplemented with 200 mM NaCl for 28 days. Histogram shows the quantification of green rosette size, $n = 3$. Student's t-test was used to test significant differences between WT and *CDEF1* line (**$P < 0.01$, *$P < 0.05$, ns, not significant). Scale bar, 1 cm. **d**, Trait contribution of individual elements to Dim1 and Dim2 from the PCA shown in Fig. 1d. Arrow length and color indicate each element's contribution to the corresponding principal component. **e**, Histogram of trait contributions to Dim1 and Dim2. The red line indicates the significance threshold above which a trait is considered a major contributor to the respective dimension.
