## Extended Figure 5 for "GWAs reveals SUBER GENE1-mediated suberization via Type One Phosphatases"

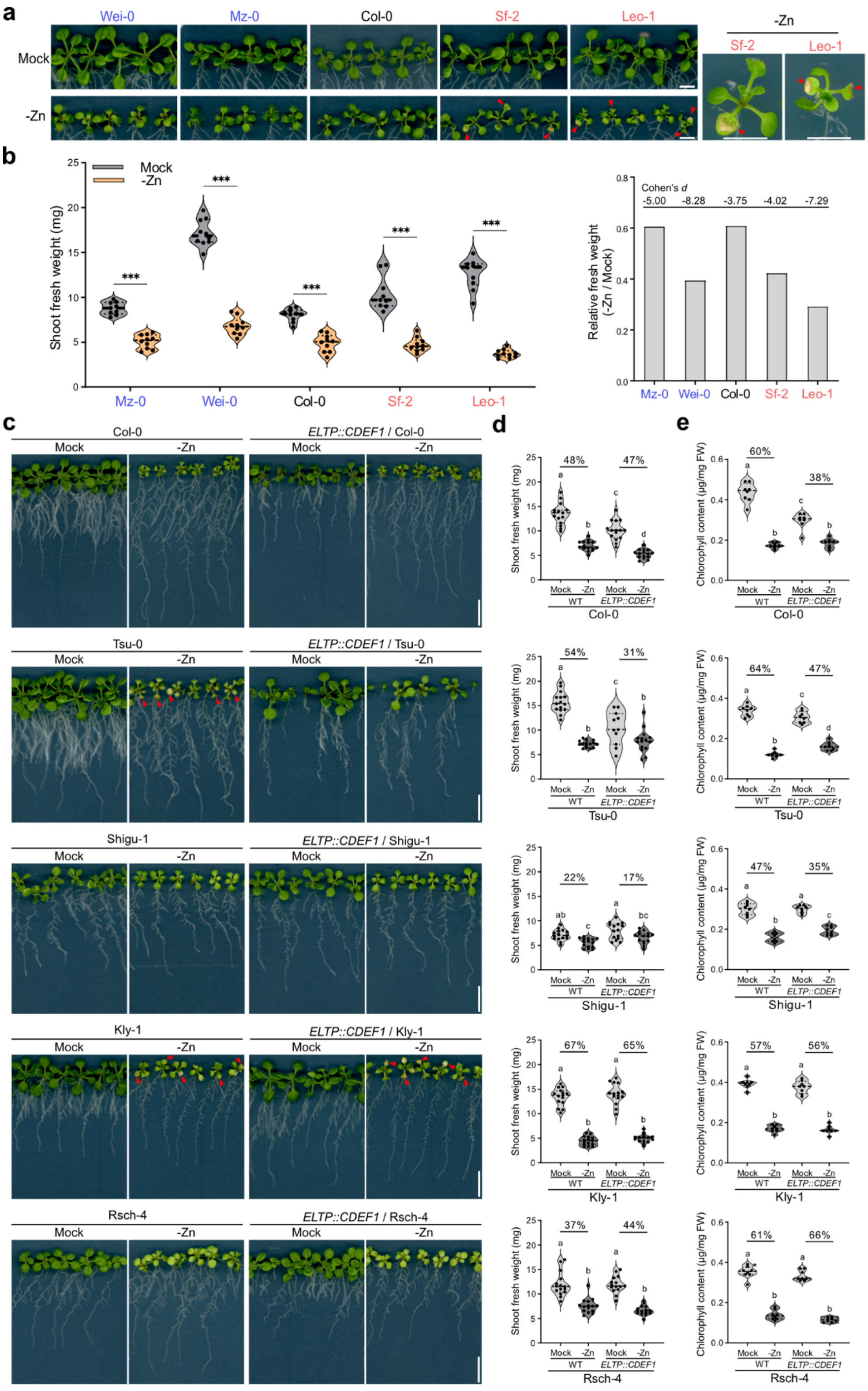


**Extended Data Fig. 5 | Effect of zinc deficiency on extreme accessions. a**, Shoot phenotype of 14-day-old extreme accessions grown under control (mock) or Zn-deficient conditions. Enlarged view of Sf-2 and Leo-1 under Zn deficiency, highlights the presence of chlorotic spots (red arrows). Scale bars, 0.5 cm. **b**, Shoot fresh weight of extreme accessions grown under mock or Zn-deficient conditions for 14 days. Data are shown as violin plots with individual values overlaid, *n* = 10. Thick dashed lines in the violin plots indicate the median, while dotted lines represent the first and third quartiles. Two-way ANOVA was performed followed by Bonferroni's test to assess significance between treatments for each accession (****P* < 0.001). Relative shoot fresh weight under mock and Zn-deficient conditions (right panel) is presented to compare the difference and Cohen's *d* values are shown on the top of histogram to evaluate the effect sizes by -Zn for each accession. **c**, 14-day-old seedlings of specified accessions without or with *ELTP::CDEF1* constructs (as described in Extended data Fig. 4a) grown under mock or -Zn conditions. Scale bar, 1 cm. Red arrows indicate chlorotic spots under -Zn. **d**, Shoot fresh weight of seedlings from (**c**). Violin plots presented as in (**b**) showing shoot fresh weight in specified genotypes and conditions, *n* = 15. One-way ANOVA followed by a Tukey's test was used to assess significant differences between genotypes; different letters indicate significant differences between conditions or genotypes (*P* < 0.05). The reduction ratios of fresh weight by -Zn condition are presented on the graphs. **e**, Chlorophyll contents of seedlings from (**c**). Data are presented and statistically analyses as in (**d**), *n* = 8.
