## Extended Figure 6 for "GWAs reveals SUBER GENE1-mediated suberization via Type One Phosphatases"

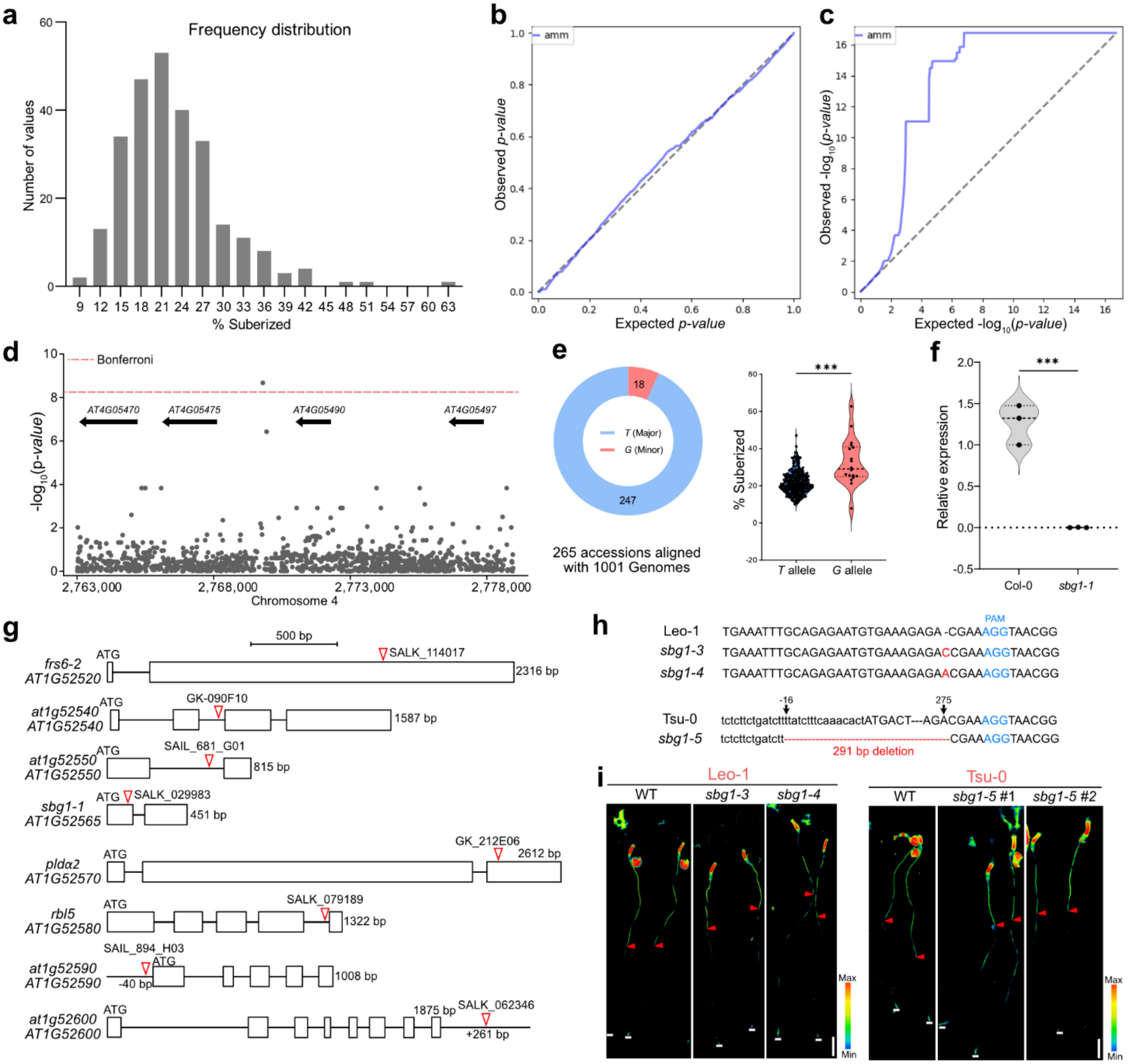


**Extended Data Fig. 6 | Mutations of *SBG1* in strong accessions reduces suberin. a**, Frequency distribution of the fully suberized zone across 265 *Arabidopsis* accessions used for the GWAS shown in Fig. 2a. **b**,**c**, Quantile-quantile (QQ) plots corresponding to the GWAS shown in the Manhattan plot in Fig. **2a**. **d**, SNPs located at positions 2,769,796 and 2,769,807 on Chromosome 4. **e**, Percentage of the suberized zone in accessions carrying the 2 alleles at SNP position 19,579,742 on Chromosome 1. Data are presented as violin plots with outlier values overlaid. Thick dashed lines in the violin plots indicate the median, while dotted lines represent the first and third quartiles. **f**, qRT-PCR analysis of *SBG1* relative transcript levels in roots of Col-0 and *sbg1-1*. Violin plots presented as in (**e**) showing the qRT-PCR results, $n$ = 3. A two-tailed Student's t-test was performed, ***$P$ < 0.001. **g**, Schematic showing the position of the T-DNA in the gene candidates for which mutants were characterized in Fig. 2c. Homozygous mutants were isolated by genotyping and T-DNA insertion sites were confirmed by sequencing. **h**, Schematic showing the position of CRISPR-generated mutations for *SBG1* in Leo-1 and Tsu-0 backgrounds. **i,** Whole-mount Fluorol Yellow-stained roots of *sbg1* mutants in Leo-1 and Tsu-0 backgrounds. Red arrows indicate the onset of the suberized zone; white lines mark the root tip. Scale bar, 1 mm.
