## Extended Figure 7 for "GWAs reveals SUBER GENE1-mediated suberization via Type One Phosphatases"

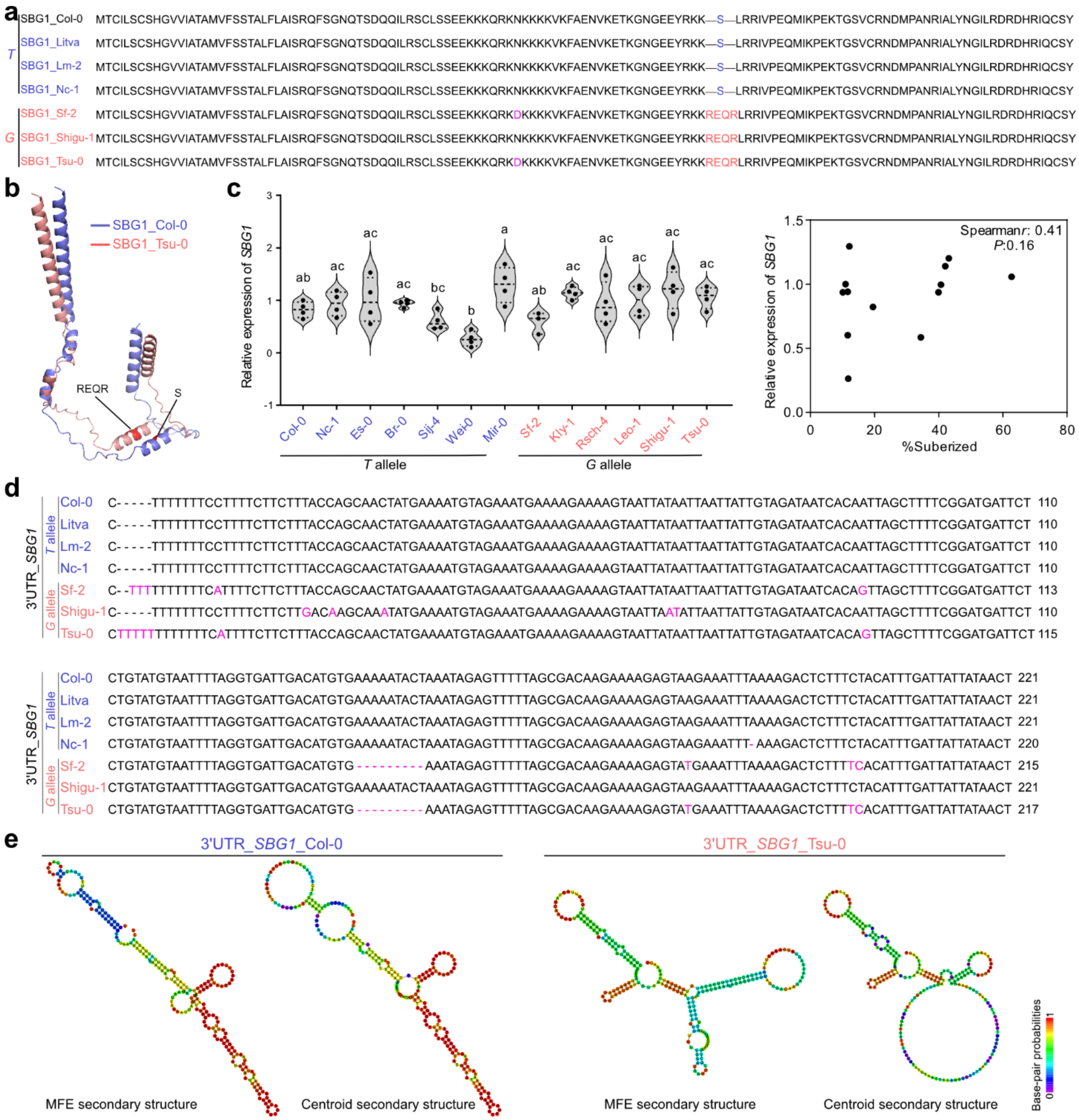


**Extended Data Fig. 7 | Alignment of protein and 3'UTR sequences of *SBG1* from *T* and *G* alleles. a**, Protein sequence alignment of SBG1 in *T* and *G* alleles. The differential sites are marked in different colors. **b**, AlphaFold predicted structures of SBG1 from Col-0 and Tsu-0. The differential sites of "REQR" and "S" were marked in red. **c**, qRT-PCR analysis of *SBG1*'s expression in the accessions with *T* and *G* alleles. Data are shown as violin plot with individual values overlaid, $n = 4$. Thick dashed lines in the violin plots indicate the median, while dotted lines represent the first and third quartiles. One-way ANOVA followed by Tukey's test was used to assess the difference and different letters indicate significant differences between accessions ($P < 0.05$). No significant correlation was shown between the expression level of *SBG1* and fully suberized pattern ($P > 0.05$). **d**, 3'UTR alignment of *SBG1* in *T* and *G* alleles. The differential sites are marked purple. **e**, RNAfold predicted RNA secondary structure of 3'UTRs of SBG1 from Col-0 (*T* allele) and Tsu-0 (*G* allele). Both MFE (minimum free energy) and centroid secondary structure are presented in color code of base-pair probabilities.
