## Extended Figure 8 for "GWAs reveals SUBER GENE1-mediated suberization via Type One Phosphatases"

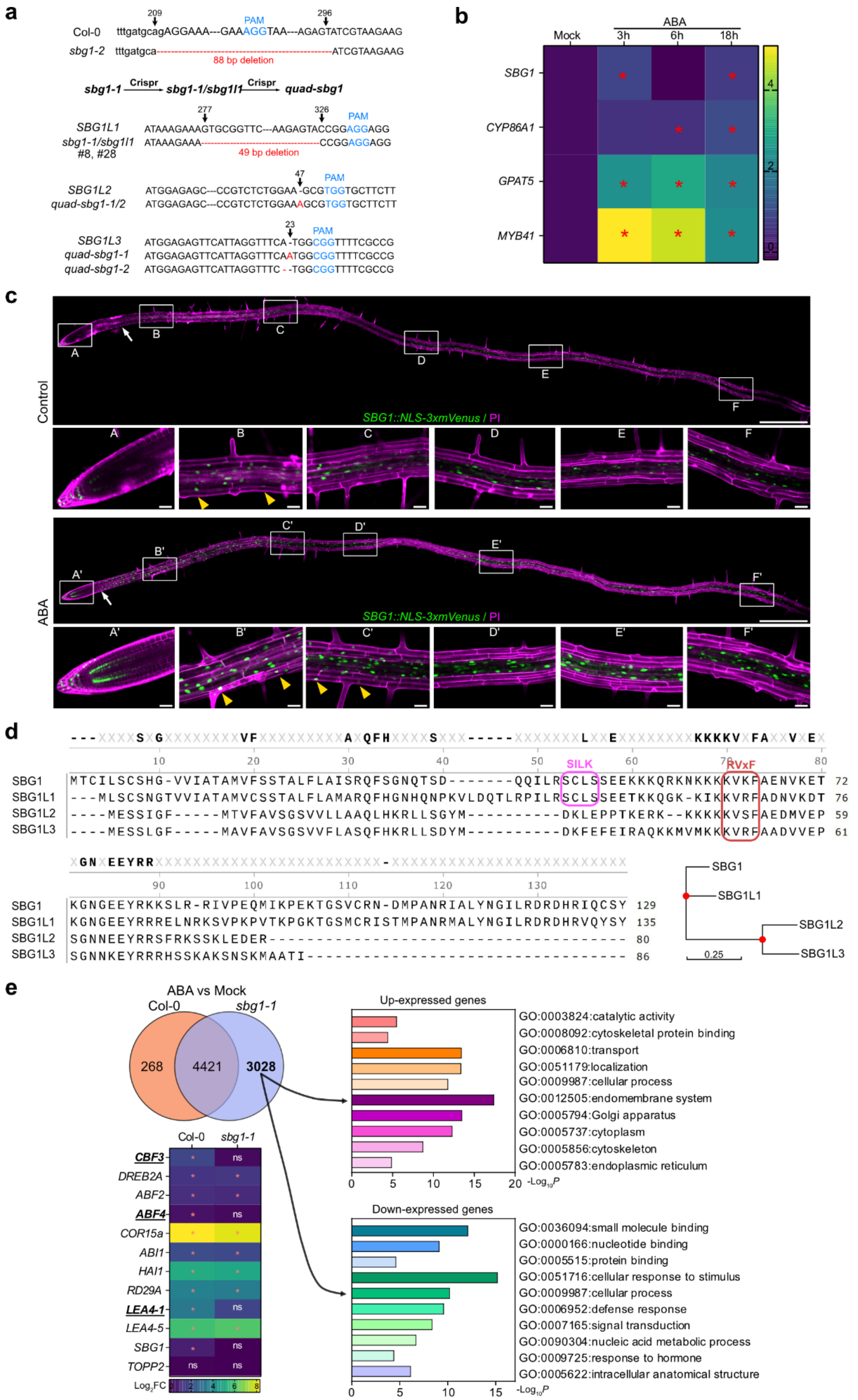

**Extended Data Fig. 8 | ABA induces the expression of *SBG1*. a**, Schematic representation of the CRISPR-generated mutation in *SBG1* and its homologs to generate the *sbg1-2*, *sbg1-1;sbg1l1* double mutant and *quad-sbg1* mutants in the Col-0 background. **b**, Heat map of qRT-PCR analysis showing relative transcript levels of *SBG1*, *CYP86A1/HORST*, *GPAT5* and *MYB41* under control (Mock) or 1 µM ABA treatment for 3, 6, or 18h. Data presented as $\log_2$(fold change) relative to mock. Student's *t*-test was used to assess significance between control and ABA treatments at each time point (**P* < 0.05, *n* = 3). **c**, Confocal images of *SBG1::NLS-3xmVenus* lines treated with or without 1 µM ABA for 18h. The full scans are identical to Fig. 4h and additional higher-resolution images for positions specified with a white box are presented. mVenus signal is shown in green and PI in magenta. White arrows indicate the onset of mVenus signal after the division zone. Yellow arrows indicate signal in the cortex or epidermis. Scale bar, 500 µm. **d**, Protein sequence alignment and phylogenetic analysis of SBG1, SBG1L1, SBG1L2 and SBG1L3. The SILK and RVxF motifs, known to mediate binding to TOPPs, are highlighted in pink and red respectively. On the phylogenetic tree, each red node represents a divergence event. **e**, Comparison of differentially expressed genes (DEG) from RNAseq in Col-0 and *sbg1-1* roots after 2 µM ABA treatment for 4h. Heat maps showing the differential expression of ABA-induced maker genes, *SBG1* and *TOPP2* in Col-0 and *sbg1-1* (**P*adj < 0.05, ns, not significant, *n* = 3 pools of approximately 250 roots). GO enrichment analysis was performed for the 3028 unique DEGs in *sbg1-1*.
