## Extended Figure 9 for "GWAs reveals SUBER GENE1-mediated suberization via Type One Phosphatases"

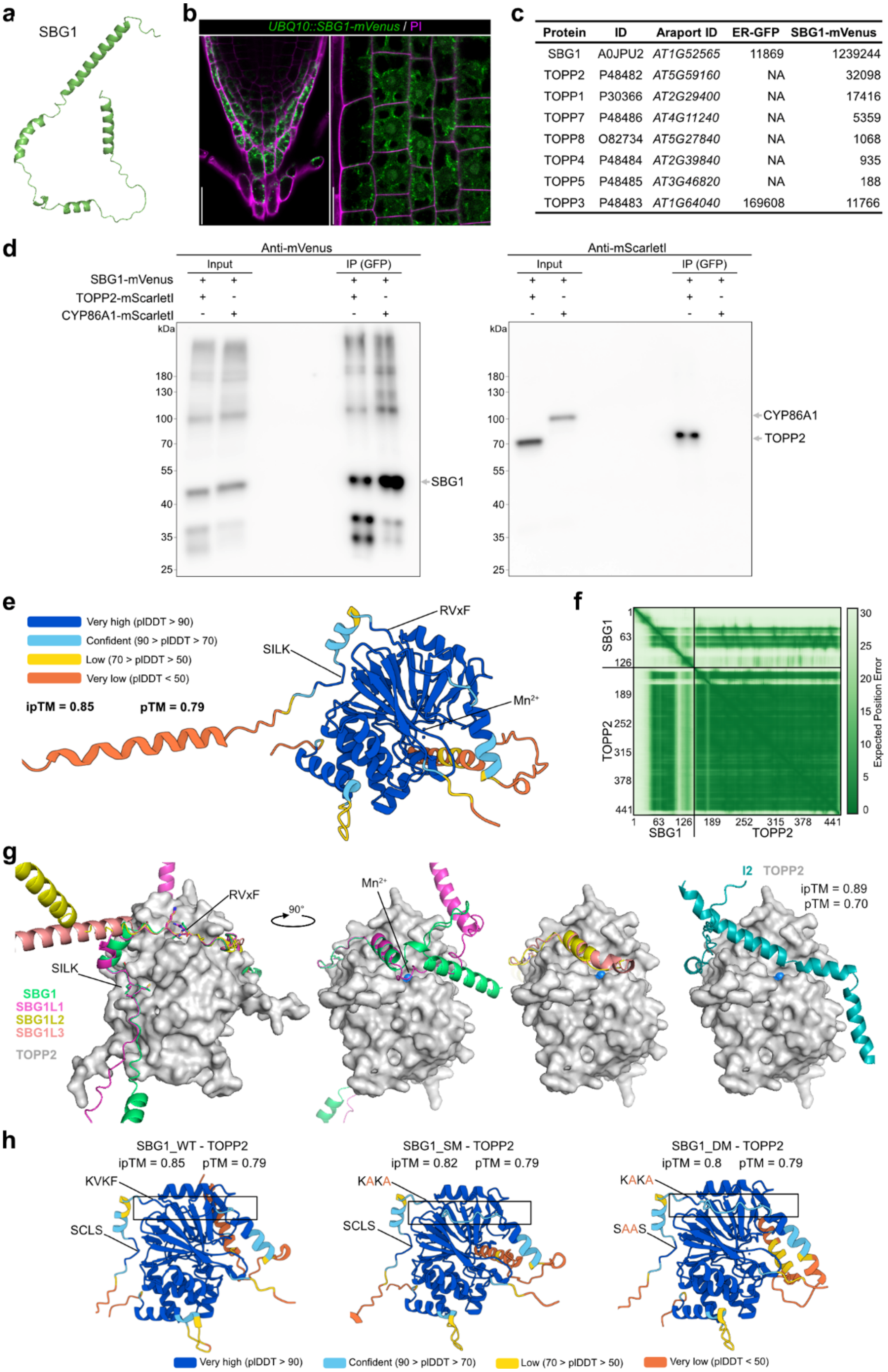


| Protein | ID | Araport ID | ER-GFP | SBG1-mVenus |
|---|---|---|---|---|
| SBG1 | A0JPU2 | *AT1G52565* | 11869 | 1239244 |
| TOPP2 | P48482 | *AT5G59160* | NA | 32098 |
| TOPP1 | P30366 | *AT2G29400* | NA | 17416 |
| TOPP7 | P48486 | *AT4G11240* | NA | 5359 |
| TOPP8 | O82734 | *AT5G27840* | NA | 1068 |
| TOPP4 | P48484 | *AT2G39840* | NA | 935 |
| TOPP5 | P48485 | *AT3G46820* | NA | 188 |
| TOPP3 | P48483 | *AT1G64040* | 169608 | 11766 |

**Extended Data Fig. 9 | Identification of TOPPs as targets of SBG1s. a**, Predicted AlphaFold structure of SBG1. **b**, Confocal images of SBG1-mVenus (green) in the *SBG1 OX4* line with PI staining (magenta) to highlight cell walls. Scale bar, 20 μm. **c**, List of TOPP candidate interactors identified by IP-MS. Peptide quantification is shown for each protein, *n* = 3. **d**, Predicted AlphaFold structure of SBG1 in complex with TOPP2. Cartoon representation colored by predicted local distance difference test (pLDDT) confidence scores. **e**, Predicted aligned error (PAE) plot corresponding to the AlphaFold model shown in (**d**). **f**, Aligned AlphaFold structures of SBG1 homologs in complex with TOPP2. Structural differences between SBG1/SBG1L1 and SBG1L2/SBG1L3 are observed at both N- and C- termini (left panels). Predicted AlphaFold structure of Inhibitor 2 (I2) in complex with TOPP2 (right panel). **g**, Full membrane views of the CoIP assay showed in Fig. 4b. **h**, AlphaFold predicted structure of SBG1_WT, SBG1_SM or SBG1_DM complex with TOPP2. Cartoon color represented as in (**d**). The mutated amino acids are marked in red; black boxes indicate the RVxF region with changed pLDDT score in SBG1_WT, SBG1_SM and SBG1_DM.
