## Extended Figure 10 for "GWAs reveals SUBER GENE1-mediated suberization via Type One Phosphatases"

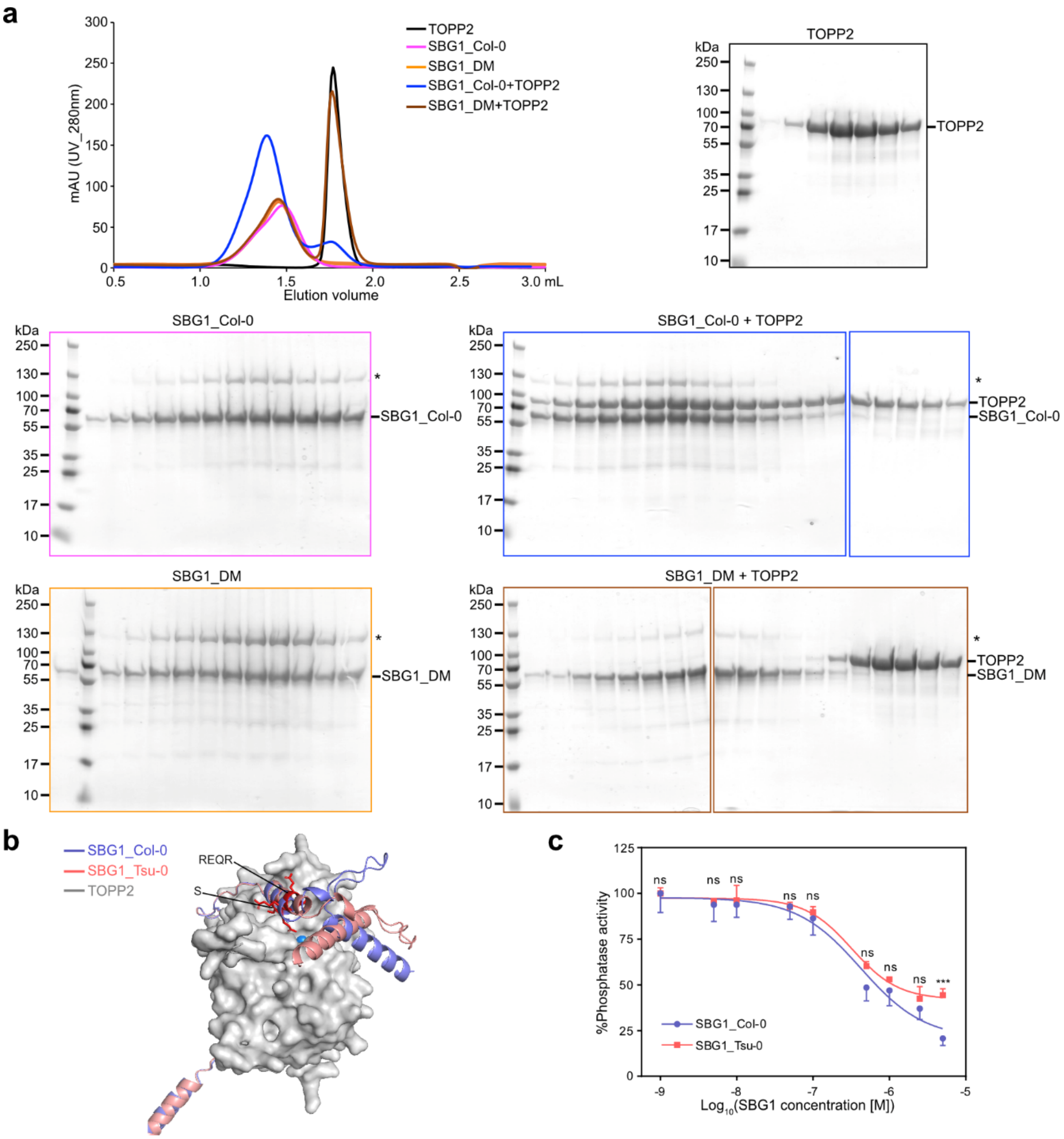


**Extended Data Fig. 10 | SBG1 binds to and inhibits TOPP2 *in vitro*.** **a**, Analytic gel-filtration assay showing that only SBG1_Col-0 but not SBG1_DM binds to TOPP2. SDS-PAGEs showing the collected fractions of eluted peaks after gel filtration. Asterisks indicate the lager band which might be the dimer complex of SBG1 in all the samples. **b**, Aligned AlphaFold predicted structure of SBG1 from Col-0 and Tsu-0 in complex with TOPP2. Side chains in the differential sites are shown in red. **c**, TOPP2 phosphatase activity assay showing comparable inhibition of TOPP2 by SBG1_Col-0 or SBG1_Tsu-0. Two-way ANOVA followed by Sidak's test was used to assess the significance at each concentration (***$P < 0.001$, ns, not significant, $n = 3$).
