## Extended Figure 11 for "GWAs reveals SUBER GENE1-mediated suberization via Type One Phosphatases"

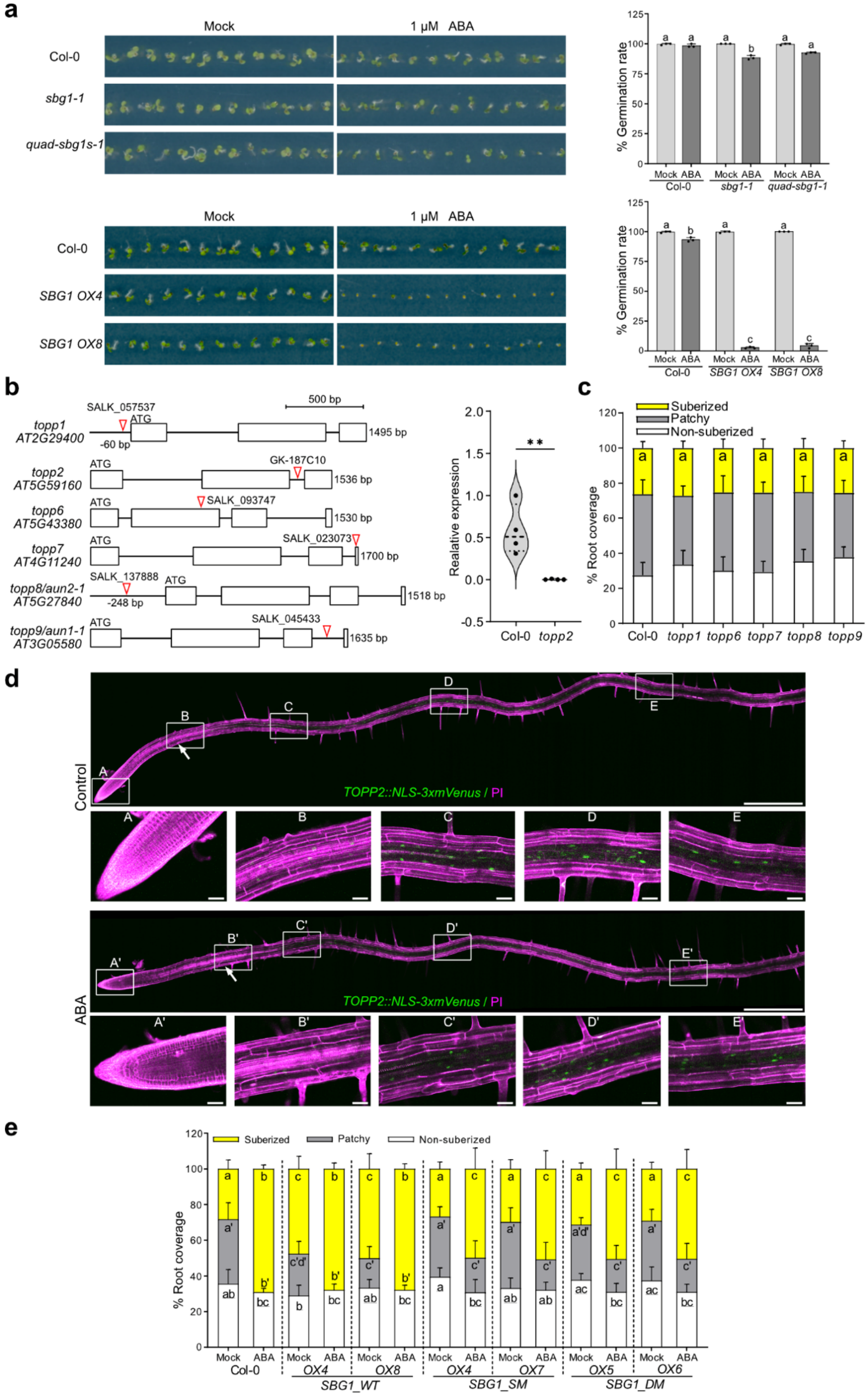


**Extended Data Fig. 11 | SBG1 promotes suberization through interaction with TOPPs.** **a**, Seed germination assay. For each replicate 84 seeds per genotypes were sown on ½ MS or ½ MS supplemented with 1 µM ABA. Germination was scored after 96 hours. Data are shown as histograms with individual values overlaid, + s.d., $n = 3$ replicates. One-way ANOVA followed by Tukey's test was used to assess differences between genotypes and different letters indicate significant differences ($P < 0.05$). **b**, Schematic showing the position of the T-DNA insertions in *topp* mutants. Homozygous mutants were isolated by genotyping and T-DNA insertion sites were confirmed by sequencing (left panel). Violin plot showing the relative expression of *TOPP2* in Col-0 and *topp2* determined by qRT-PCR (right panel). Thick dashed lines in the violin plots indicate the median, while dotted lines represent the first and third quartiles. Student's t-test was used to test the significance (**$P < 0.01$, $n = 4$). **c**, Suberin deposition pattern in *topp* mutants. Data are presented as stacked columns showing the mean percentage of each suberization zone + s.d., $n = 10$. One-way ANOVA followed by a Tukey's test was used for the suberized zone to compare genotype. No significant difference was detected ($P > 0.05$). **d**, Confocal images of *TOPP2::NLS-3xmVenus* lines treated with or without 1 µM ABA for 18h. The full scan as well as higher resolution images are presented. mVenus signal is shown in green and PI in magenta. White arrows indicate the onset of mVenus signal. Scale bar, 500 µm. **e**, Suberin deposition pattern in transgenic lines overexpressing *SBG1_WT, SBG1_SM or SBG1_DM* under control (mock) or 1 µM ABA treatment for 16 h. 4-day-old seedlings were transferred to either ½ MS plates or ½ MS plate + ABA. Data are presented as in (**c**). Different letters indicate significant differences in the different suberized zones between genotypes and conditions with apostrophes and underlines for the patch and non-suberized zones respectively.
