## Extended Figure 12 for "GWAs reveals SUBER GENE1-mediated suberization via Type One Phosphatases"

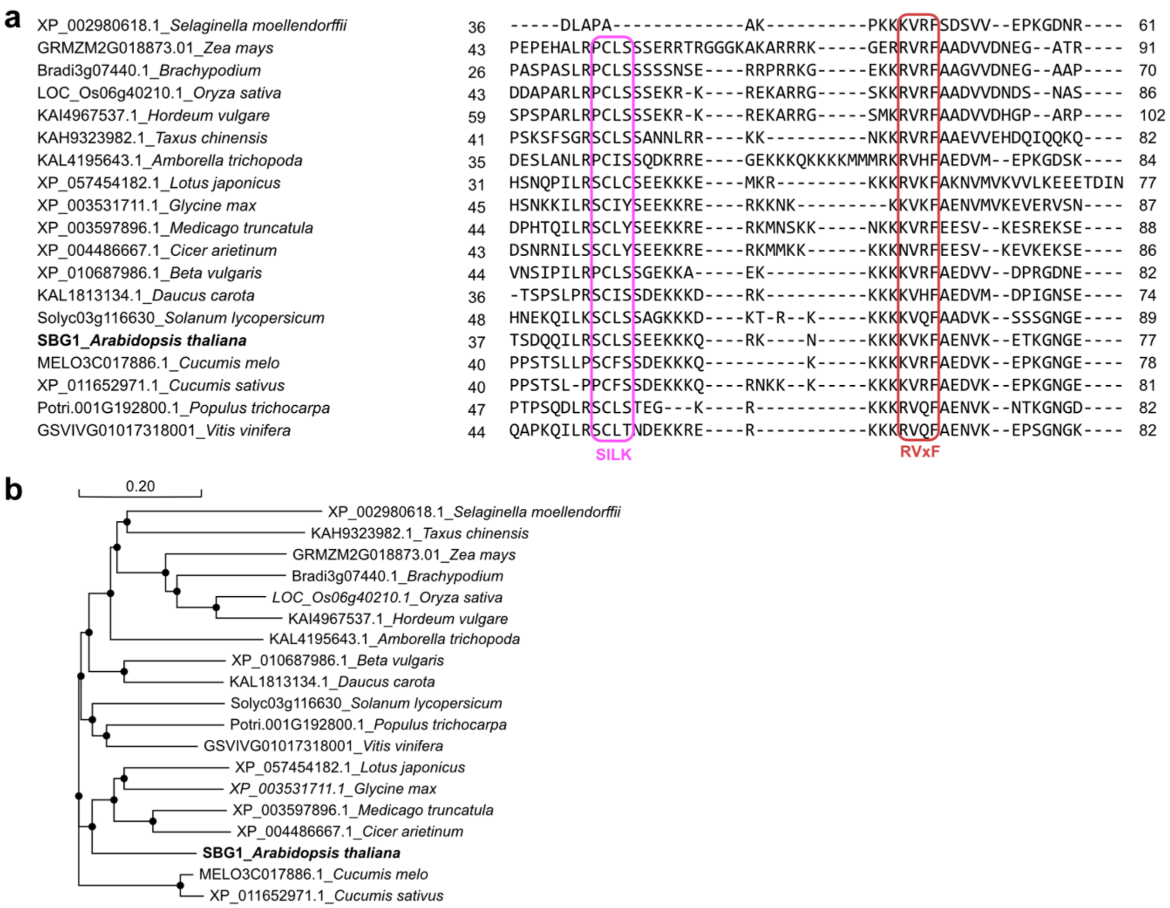


**Extended Data Fig. 12 | Phylogenetic analysis of SBG1 and orthologs across species. a**, Protein sequence alignment of SBG1 from *A. thaliana* and its orthologs from 18 plant species. The sequences of SBG1's orthologs were obtained by NCBI blast with highest protein similarity for each species. The SILK and RVxF motifs are highlighted in pink and red respectively. **b**, Phylogenetic analysis of SBG1 and its 18 orthologs. Scale bar indicates the branch lengths.
